# TooT-PLM-P2S: Incorporating Secondary Structure Information into Protein Language Models

**DOI:** 10.1101/2024.08.13.607781

**Authors:** Hamed Ghazikhani, Gregory Butler

## Abstract

In bioinformatics, modeling the protein space to better predict function and structure has benefitted from Protein Language Models (PLMs). Their basis is the protein’s amino acid sequence and self-supervised learning. Ankh is a prime example of such a PLM. While there has been some recent work on integrating structure with a PLM to enhance predictive performance, to date there has been no work on integrating secondary structure rather than three-dimensional structure. Here we present TooT-PLM-P2S that begins with the Ankh model pre-trained on 45 million proteins using self-supervised learning. TooT-PLM-P2S builds upon the Ankh model by initially using its pre-trained encoder and decoder. It then undergoes an additional training phase with approximately 10,000 proteins and their corresponding secondary structures. This retraining process modifies the encoder and decoder, resulting in the creation of TooT-PLM-P2S. We then assess the impact of integrating secondary structure information into the Ankh model by comparing Ankh and TooT-PLM-P2S on eight downstream tasks including fluorescence and solubility prediction, sub-cellular localization, and membrane protein classification. For both Ankh and TooT-PLM-P2S the downstream tasks required task-specific training. Few of the results showed statistically significant differences. Ankh outperformed on three of the eight tasks, TooT-PLM-P2S did not outperform on any task for the primary metric. TooT-PLM-P2S did outperform for the precision metric for the task of discriminating membrane proteins from non-membrane proteins. This study requires future work with expanded datasets and refined integration methods.

## 1. Introduction

### 1.1. Background and Motivation

Protein Language Models (PLMs) are computational tools that predict protein properties and functions based on amino acid sequences [1–3]. They utilize techniques from natural language processing (NLP) to analyze primary structures, which are the linear sequences of amino acids [4–7]. PLMs have applications in predicting protein functions, interactions, and structural properties [1,3,8–14]. They capture patterns and relationships from large protein sequence datasets that are challenging to identify manually.

Recent studies have explored the integration of three-dimensional (3D) structural data into PLMs to enhance predictive performance. For instance, Heinzinger et al. utilized 3D structural information in their ProstT5 model to improve protein function prediction tasks [15]. Similarly, Wang et al. introduced S-PLM, which employs multi-view contrastive learning to align sequence and 3D structural data, enhancing clustering and classification performance [16]. These approaches leverage data from comprehensive databases like AlphaFoldDB [17] to provide additional structural context.

PLMs are valuable in various bioinformatics tasks such as predicting protein-protein interactions, annotating protein functions, and identifying functional domains [1,3,8–10]. They also aid in understanding the effects of mutations, crucial for protein engineering and drug development [9,10,18]. One significant advancement is the Ankh model, which enhances protein function prediction by incorporating features from primary amino acid sequences and employing advanced architectural and training strategies to achieve high efficiency with fewer parameters [19].

Protein secondary structures, such as alpha-helices and beta-sheets, are local folded shapes within polypeptides resulting from amino acid interactions [20,21]. These structures are critical for determining the overall three-dimensional shape and function of proteins [22]. Understanding secondary structures is essential for predicting protein folding, interactions, and biological roles [23].

Integrating secondary structure information into PLMs may enhance their predictive capabilities by providing additional context beyond the primary sequence. Secondary structures offer insights into spatial arrangements of amino acids, important for protein stability, dynamics, and interactions. Incorporating this data could lead to more accurate predictions of protein functions and behaviors.

### 1.2. Literature Review

Recent advancements in Protein Language Models (PLMs) have significantly improved our understanding of protein sequences and their functionalities [1,3,11–14,19]. Notable models such as ProstT5 [15] and S-PLM [16] integrate both sequence and three-dimensional (3D) structure data to enhance protein function predictions. ProstT5 utilizes a combination of amino acid sequences and 3D structural information encoded as token sequences, leveraging data from AlphaFoldDB [17] to improve prediction tasks and generate novel protein sequences with specific structural scaffolds. Similarly, S-PLM employs multi-view contrastive learning to align sequence and 3D structural information within a coordinated latent space, improving performance in clustering and classification tasks.

Despite these advancements, integrating secondary structure information into PLMs remains a critical gap. Secondary structures, such as alpha-helices and beta-sheets, represent local folded shapes within a polypeptide that determine the overall three-dimensional conformation and functional properties of proteins. Secondary structure information can significantly enhance the accuracy of protein folding predictions, functional site identification, and interaction interface modeling.

Many existing models focus primarily on the primary sequence of amino acids without adequately incorporating the structural context provided by secondary structures. This limitation restricts the models’ predictive accuracy and their ability to generalize across diverse protein datasets. Models like SES-Adapter [24] and SaProt [25] have attempted to address this by incorporating structure-aware vocabularies and adapter modules, but the potential of integrating secondary structure information remains underexplored.

Our research aims to address this gap by developing a PLM that explicitly integrates secondary structure information into its training and prediction processes. This approach aims to enhance the model’s predictive accuracy and provide a more comprehensive understanding of protein functionalities, particularly for tasks involving membrane proteins and other complex protein structures.

### 1.3. Research Problem and Objectives

The current generation of Protein Language Models (PLMs) primarily focuses on the primary structure of proteins, the linear sequence of amino acids, often neglecting secondary structures like alpha-helices and beta-sheets. These secondary structures are crucial for determining the three-dimensional conformation and functional properties of proteins. The lack of secondary structure information limits the predictive accuracy and generalizability of PLMs in tasks such as protein classification, function prediction, and interaction modeling.

The objective of this research is to evaluate the effectiveness of integrating secondary structure information into a pretrained PLM. Specifically, we aim to determine whether this integration enhances the model’s performance across various protein-related tasks. The integration was achieved using an encoder-decoder architecture based on the Ankh model, where the primary amino acid sequence is input to the model, and the model generates the corresponding secondary structure. This approach implicitly integrates secondary structure knowledge into the pretrained model, which has been extensively trained on primary sequence data.

By incorporating secondary structure data, we hypothesize that the model will achieve a more comprehensive understanding of protein behavior and interactions. The enhanced model, named TooT-PLM-P2S, was assessed across multiple tasks using diverse datasets. This evaluation includes fluorescence prediction, solubility prediction, sub-cellular localization prediction, ion channel classification, transporter classification, membrane protein classification, and secondary structure prediction. We also conducted a detailed analysis of misclassification cases to understand the underlying reasons for errors, employing three distinct bioinformatics-based methods: Multiple Sequence Alignment (T-Coffee), Orthologous Groups (eggNOG) Analysis, and Motif Alignment and Search Tool (MEME Suite).

The primary objectives of this study are as follows:

1. Integration of Secondary Structure Information: Develop an enhanced version of the Ankh model by incorporating secondary structure information into the training process. This aims to provide the model with an intermediate understanding of the protein’s three-dimensional conformation, leading to more accurate protein classification and function prediction.
2. Evaluation of Classification Accuracy: Rigorously evaluate the classification performance of TooT-PLM-P2S in comparison to the baseline Ankh model. This involves diverse tasks and datasets, including fluorescence, solubility, localization, ion channels, transporters, membrane proteins, and secondary structure prediction, to assess the impact of secondary structure integration on classification accuracy.
3. Analysis of Failure Cases: Conduct a detailed analysis of instances where both the baseline and TooT-PLM-P2S models misclassify protein sequences. This analysis will use tools such as Multiple Sequence Alignment (T-Coffee), Orthologous Groups (eggNOG) Analysis, and Motif Alignment and Search Tool (MEME Suite) to uncover patterns or specific characteristics that lead to misclassifications, providing insights into the limitations and areas for improvement in PLMs.

This structured approach ensures a comprehensive evaluation of the TooT-PLM-P2S model, providing insights into the benefits of integrating secondary structure information into PLMs.

### 1.4. Contribution of This Paper

This paper makes several key contributions to the field of protein language modeling and bioinformatics: This study presents the development of a protein language model, TooT-PLM-P2S, which integrates secondary structure information into the training process, building upon the existing Ankh model [19]. By incorporating secondary structure information, TooT-PLM-P2S demonstrates improved predictive metrics for certain protein function and classification tasks, such as fluorescence prediction.

The paper provides a comprehensive comparison between TooT-PLM-P2S and the baseline Ankh model, benchmarking performance across multiple downstream tasks. This highlights the benefits of integrating secondary structure information.

We also conduct an enrichment analysis of misclassifications made by both models using tools such as Multiple Sequence Alignment (T-Coffee), Orthologous Groups (eggNOG) Analysis, and Motif Alignment and Search Tool (MEME Suite). This analysis identifies patterns and characteristics that lead to misclassifications, offering crucial insights for understanding model limitations and guiding future improvements.

### 1.5. Structure of the Paper

This paper is organized into several sections. Section 2 outlines the methodological framework used in the study, detailing the model architecture, integration of secondary structure knowledge, and the evaluation approach. Section 3 presents the findings from the evaluation of the TooT-PLM-P2S model. Section 4 discusses the implications of these results. Section 5 summarizes the key outcomes of the study, highlighting the contributions and potential future directions for research in protein language modeling.

## 2. Materials and Methods

### 2.1. Model Architecture: TooT-PLM-P2S

To enhance protein language models (PLMs), we developed the TooT-PLM-P2S model by continuing the training of the Ankh model using a secondary structure dataset. This process does not modify the Ankh architecture; it simply extends its training to include secondary structure information. The Ankh model remains intact throughout this process. For downstream tasks, we used ConvBERT with frozen representations from these models without further fine-tuning, following methodologies from recent studies [3,26,27]. This approach allows us to assess the predictive capabilities of the representations learned by TooT-PLM-P2S in various protein-related tasks.

#### Ankh Model Overview

The Ankh model [19] is a highly efficient PLM with significantly fewer parameters compared to traditional models. It uses Google’s TPU-v4 technology, achieving state-of-the-art results with fewer than 10% of the parameters typically required for pre-training and less than 7% for inference tasks. This efficiency makes the model accessible for broader research applications. Ankh is available in two versions: Ankh base with 726 million parameters and Ankh large with 1.9 billion parameters. In our experiments, we used the Ankh base model due to its smaller size, which suited the available resources.

The Ankh model excels in various protein modeling tasks, such as structure prediction and functional benchmarking. It is effective in generating protein variants using insights into evolutionary conservation and mutation trends, supporting complex biological research. The architecture includes 48 encoder layers and 24 decoder layers, utilizing a Gated-GELU activation function and a relative positional embedding dimension of 32 with an offset of 128. Pre-trained on the UniRef50 dataset, Ankh demonstrates robustness and accuracy across different protein-related tasks.

Pre-training Ankh involved masked language modeling, where 20% of the input tokens were masked, and the model was trained to predict these masked tokens. This approach allows the model to learn underlying patterns and relationships within protein sequences. The embedding dimension is set at 768, with 12 attention heads, capturing both local and global sequence features effectively. Training was optimized using the Adafactor optimizer and a linear scheduler, with a learning rate of 0.004 and a batch size of 24. This training strategy enables Ankh to achieve high performance in protein modeling tasks with a significantly reduced computational burden.

### 2.2. Integration of Secondary Structure Knowledge

To develop TooT-PLM-P2S, we extended the Ankh model by incorporating secondary structures like alpha-helices and beta-sheets. This integration aims to improve the model’s predictive accuracy and understanding of protein functions.

The integration leverages the Ankh model’s sequence-to-sequence architecture, specifically adopting its 48 encoder layers and 24 decoder layers. We used the initial weights from the pre-trained Ankh model, which had been pre-trained on 45 million primary protein sequences. This provided a strong foundation of learned representations.

Primary protein sequences are input into the model, which predicts their corresponding secondary structures. The encoder processes the primary sequence and creates a contextual representation of each amino acid. This representation is passed to the decoder, which predicts the secondary structure for each amino acid. A cross-entropy loss function measures the model’s performance by quantifying the difference between the predicted and actual distributions of secondary structure types at each position.

To create the TooT-PLM-P2S model (Figure 2), we further trained the Ankh model using a dataset that pairs primary protein sequences with their annotated secondary structures. Training was performed over 10 epochs, determined through hyperparameter optimization with Optuna, to minimize the discrepancy between predicted and actual secondary structures.

**Figure 1.**
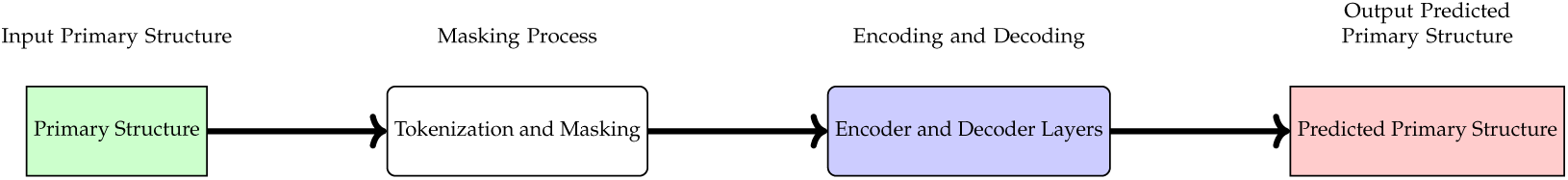
Ankh model architecture. The plot of the Ankh model architecture showing primary structure input, tokenization and masking, encoding-decoding process, and predicted primary structure output.

**Figure 2.**
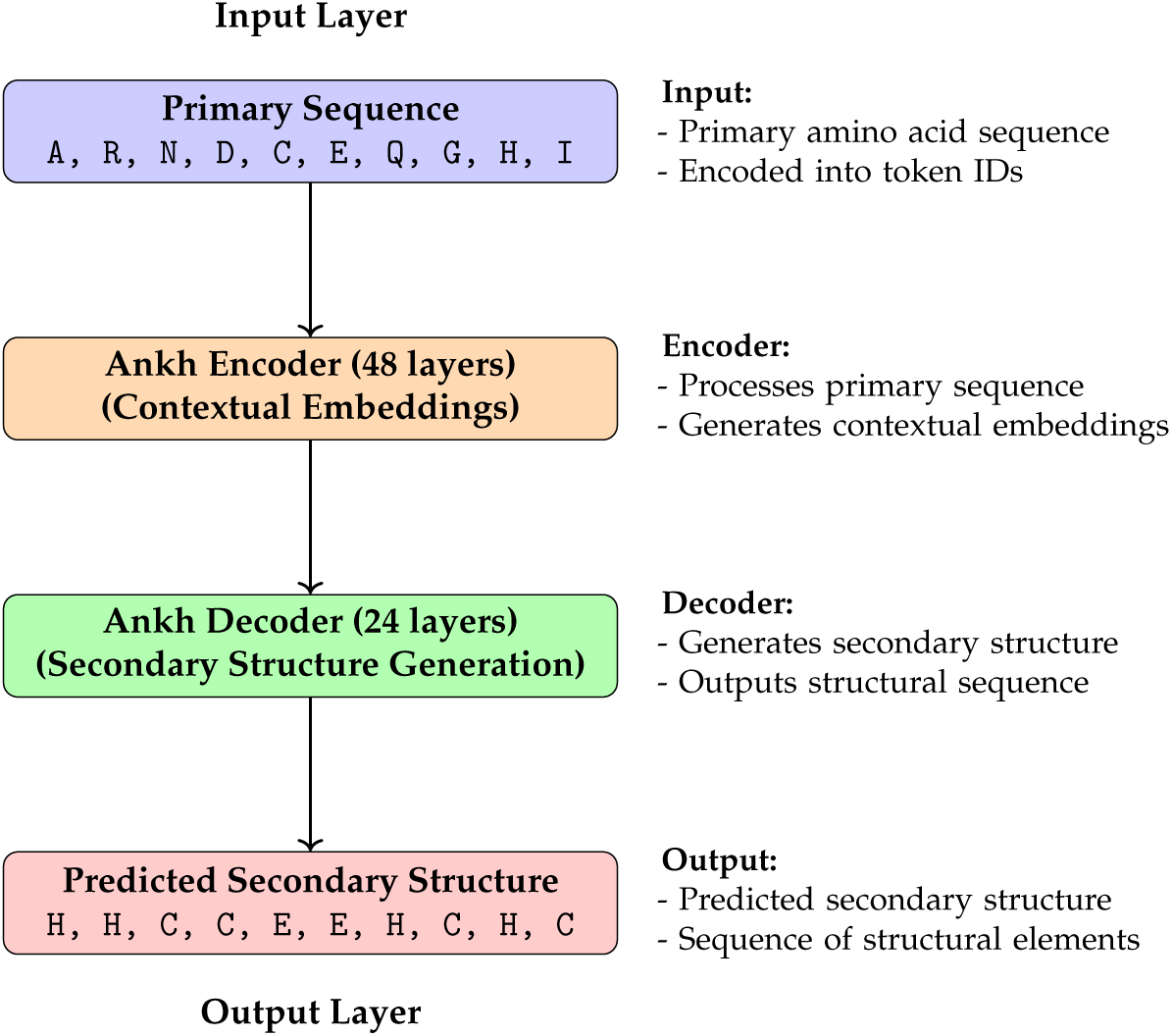
Schematic of the TooT-PLM-P2S Model Architecture. The model integrates secondary structure information to enhance the predictive capabilities of the protein language model. For the training the three-state of secondary structure has been used.

Optuna was used to fine-tune the following parameters: learning rate (0.0003), training epochs (10), warmup ratio (0.2), weight decay (0.09), gradient accumulation steps (4), and batch size (1). The training dataset was sourced from NetSurfP-2.0 [28], designed for secondary structure prediction. It includes 11,361 protein sequences, with 8,678 training samples, 2,170 validation samples, and 513 test samples. This dataset provides classifications at both 3-class and 8-class levels, ensuring rigorous evaluation and facilitating direct comparison with the baseline Ankh model. The training used the three-state secondary structure classification.

### 2.3. Downstream Tasks

This section details the prediction tasks used to evaluate TooT-PLM-P2S, examining various aspects of the model’s capabilities. These tasks employ diverse datasets, reflecting real-world challenges and benchmarks used by PEER [29] and xTrimoPGLM [30], as well as our prior projects.

We selected these tasks to span a wide range of protein functionalities and structures, ensuring comprehensive evaluation. This includes datasets focusing on membrane proteins, ion channels, and transporters, emphasizing the importance of these categories. Three specific membrane protein datasets were included to highlight membrane protein classification’s significance. This broad selection ensures that TooT-PLM-P2S’s performance is rigorously tested across various protein function prediction tasks, demonstrating the model’s versatility and robustness.

#### 2.3.1. Non-SSP Tasks

##### Fluorescence Prediction (FluP)

This regression task assesses the fluorescence intensity of green fluorescent protein mutants, which is crucial for tracking proteins in live cells and organisms. The dataset, annotated by Sarkisyan et al. [31], includes mutants with up to three mutations for training and evaluation, and mutants with four or more mutations for testing. This setup tests the model’s ability to generalize from lower-order to higher-order mutations.

##### Solubility Prediction (SolP)

Solubility prediction is vital for designing effective pharmaceuticals, as soluble proteins are more likely to be functional and usable in drug formulations. This binary classification task uses the DeepSol dataset [32], ensuring no protein in the training and evaluation sets shares more than 30% sequence identity with any protein in the test set to prevent information leakage.

##### Sub-cellular Localization Prediction (LocP)

Predicting protein localization within the cell is important for understanding protein function and interactions, especially in disease research. The DeepLoc dataset [33] classifies proteins into 10 sub-cellular localizations. This dataset is critical for evaluating the model’s capability to capture functional context from protein sequences.

##### Ion Channels Prediction (IonP)

Differentiating ion channels from other membrane proteins is essential for understanding various physiological processes and drug target identification. The dataset used is from the DeepIon project [34], compiled from UniProt and refined to reduce sequence similarity below 20%. It includes ion channels, ion transporters, and other membrane proteins, with ion transporters excluded for consistency with previous methodologies.

##### Transporters Prediction (TranP)

Identifying transporters is crucial for studying how substances move across cellular membranes, which is fundamental in cellular biology and pharmacology. This task uses a dataset from the TrSSP project [35], consisting of well-characterized transporter, carrier, and channel proteins from SwissProt. Sequences with fragmented or ambiguous annotations are excluded to ensure data quality.

##### Membrane Proteins Prediction (MemP)

Differentiating membrane proteins from other proteins helps in understanding cellular processes and the role of membrane-bound proteins in signaling and transport. This dataset is the same as that used in the TooT-M project [36], derived from Swiss-Prot. It filters for sequence quality and diversity, excluding sequences with inferred homology, less than 50 amino acids, lacking molecular function annotations, or exhibiting over 60% pairwise similarity.

#### 2.3.2. SSP Task

##### SSP3 (3-State Secondary Structure Prediction)

SSP3 involves predicting three states of secondary structure (alpha-helices, beta-sheets, and coils), which are fundamental for understanding protein folding and function. This task uses the NetSurfP-2.0 [28] dataset. Additional testing sets include CB513 [37], TS115 [38], CASP12 [39], and CASP14 [40].

##### SSP8 (8-State Secondary Structure Prediction)

SSP8 involves predicting eight distinct states of secondary structure, providing a more detailed and nuanced understanding of protein folding patterns. This task also uses the NetSurfP-2.0 dataset with the same additional testing sets (CB513, TS115, CASP12, and CASP14) to ensure robustness.

By addressing these diverse tasks, the TooT-PLM-P2S model is comprehensively evaluated, demonstrating its versatility and robustness across various aspects of protein function prediction.

#### 2.3.3. Overview of ConvBERT Use

ConvBERT [41] (Figure 3) is employed as the downstream model in our architecture for protein structure prediction tasks due to its effectiveness in managing complex sequences. ConvBERT integrates convolutional layers with self-attention mechanisms, capturing both local and global sequence features. For each prediction task, the pre-trained TooT-PLM-P2S model generates frozen embeddings, which are then fed into ConvBERT for final predictions. This approach retains the benefits of the pre-trained models without altering their parameters, ensuring consistency across tasks.

**Figure 3.**
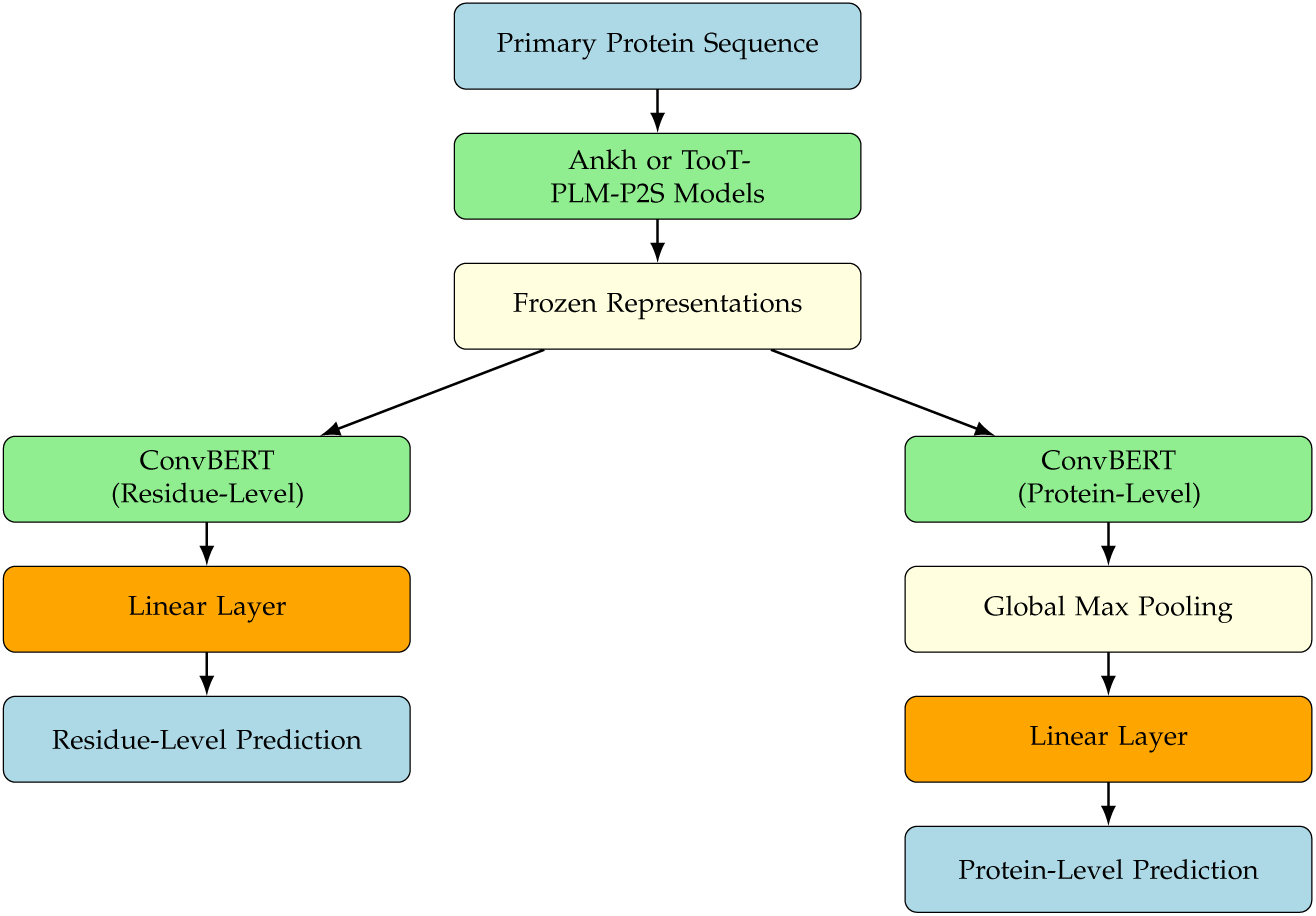
Downstream tasks using ConvBERT. This plot illustrates the process of applying the TooT-PLM-P2S or Ankh models for various downstream tasks.

The ConvBERT configuration used includes an embedding dimension that matches the pre-trained model, a feed-forward network dimension set to half the embedding dimension, four attention heads, a dropout rate of 0.2, a convolutional kernel size of 7, and the GatedGELU activation function [42]. The ConvBERT classifier comprises a ConvBERT layer followed by linear layers tailored to the task type, with no activation for regression tasks, sigmoid for binary classification, and softmax for multi-class classification. A global max pooling layer aggregates features from the convolutional layer outputs for classification.

Hyperparameter tuning for ConvBERT is conducted using Optuna [43], focusing on optimizing parameters such as learning rate, weight decay, warmup ratio, and gradient accumulation steps. The goal is to maximize the Matthews Correlation Coefficient (MCC) on the validation set. MCC is chosen for its robustness in evaluating classifier performance, particularly with imbalanced datasets [44].

### 2.4. Model Evaluation and Validation

#### 2.4.1. Evaluation Metrics

To comprehensively assess the performance of the TooT-PLM-P2S model across various prediction tasks, we employed a set of well-established evaluation metrics, each tailored to the specific nature of the tasks. These metrics provided a detailed understanding of the model’s predictive capabilities and facilitated robust comparisons with the baseline Ankh model.

1. Accuracy:

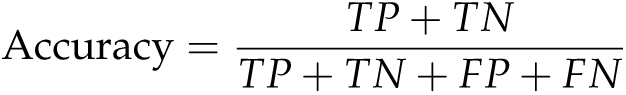

Accuracy measures the proportion of true results (both true positives and true negatives) among the total number of cases examined. It is particularly useful for balanced datasets.

2. Precision:

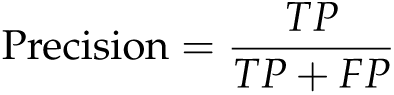

Precision, also known as positive predictive value, calculates the fraction of relevant instances among the retrieved instances.

3. Recall:

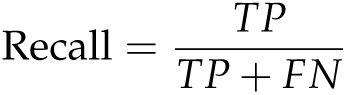

Recall, or sensitivity, measures the ability of the model to identify all relevant instances.

4. F1 Score (F1):

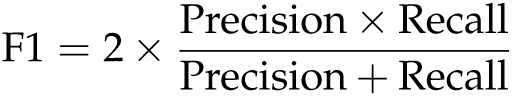

The F1 score is the harmonic mean of precision and recall, providing a single metric that balances both concerns.

5. Matthews Correlation Coefficient (MCC):

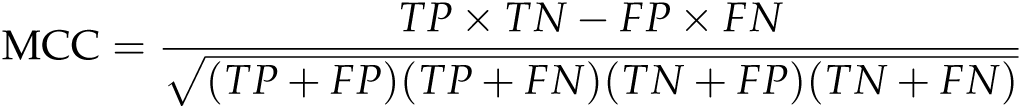

MCC is particularly informative for imbalanced datasets as it considers true and false positives and negatives.

6. Spearman’s *ρ* Correlation:

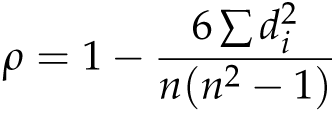

Spearman’s *ρ* measures the rank correlation between predicted and actual values, used in regression tasks to assess how well the model predicts the ordering of data points. Here, *d_i_*represents the difference between the ranks of the *i*-th pair of values and *n* is the total number of data points.

These metrics were selected for their relevance to the specific nature of each prediction task, ensuring an evaluation of both classification and regression models.

#### 2.4.2. Validation Techniques

To ensure robust validation of the TooT-PLM-P2S model’s performance, we employed multiple evaluation techniques.

##### 5-Fold Cross-Validation

We used 5-Fold Cross-Validation to provide a thorough assessment of the model’s generalizability and to mitigate the risk of overfitting. In this approach, the dataset for each task was divided into five equal parts. Each part was used once as a validation set while the remaining four parts formed the training set. This process was repeated five times, allowing every data point to be used for both training and validation. Cross-validation ensures that the model’s performance is evaluated across different subsets of the data, providing a more reliable and generalized performance metric. The best hyperparameters, identified using Optuna, were consistently applied across all folds to ensure comparability.

##### Independent Test Set Evaluation

Following cross-validation, we evaluated the model’s generalization capabilities on an independent test set, distinct from the training and validation sets. This step was crucial for assessing the real-world applicability of the model. By using a completely separate dataset for this evaluation, we ensured that the performance metrics reflected the model’s ability to generalize beyond the data it was trained and validated on.

##### Statistical Tests

To determine the statistical significance of the performance differences between the TooT-PLM-P2S and the baseline Ankh models, we conducted t-tests [45] on the cross-validation results. The t-tests helped verify that the observed differences in performance metrics were not due to random variations but were statistically significant. We used a threshold of 5*e^−^*^2^ for determining significance. This rigorous statistical analysis ensured that the improvements or changes in model performance were genuine and not attributable to chance.

### 2.5. Enrichment Analysis of Misclassifications

To gain deeper insights into the misclassifications made by our models, we employed several analytical tools focusing on sequence alignment, functional annotation, and motif identification. Misclassifications offer insights into the limitations of a model and highlight areas for iterative improvement, ensuring the model’s applicability across a wide range of proteins with varying complexities. In this context, an Enrichment Analysis (EA) [46,47] was conducted on the misclassified sequences by both the baseline Ankh model and the newly developed TooT-PLM-P2S model. This analysis utilized a suite of bioinformatics tools—T-Coffee, eggNOG, and the Motif Alignment and Search Tool (MAST)—each providing unique perspectives on the misclassified sequences. Understanding the root causes of misclassifications is critical for improving model accuracy and reliability. Previous work [46,47] has shown that analyzing misclassified instances can uncover underlying patterns and biases in model training data.

#### 2.5.1. Sequence Alignment: T-Coffee

We used T-Coffee [48] for multiple sequence alignment (MSA) to compare misclassified sequences with correctly classified ones, aiming to identify sequence regions or motifs problematic for the models. Tasks were defined for transporters, localization, solubility, ion channels, and membrane proteins. Due to the large number of alignment files, we randomly selected five alignment files per task. Protein sequences in FASTA files were parsed using Biopython [49]. For each task, alignment files were processed to separate misclassified sequences from correctly classified ones based on sequence identifiers.

Sequence identity was calculated by comparing amino acid residues at each position, computing the proportion of matches over the alignment length. For each misclassified sequence, pairwise sequence identities were calculated with all correctly classified sequences in the same alignment file. The average sequence identity for each alignment file was computed and aggregated to obtain task-specific average identities, which were stored in a dictionary for further analysis.

#### 2.5.2. Functional Annotation: eggNOG

We used eggNOG [50] to explore the evolutionary relationships and functional contexts of misclassified sequences. By analyzing annotated data from eggNOG mapper output files, we examined the distribution of COG categories across various tasks, comparing the frequencies of orthologous groups (OGs) between misclassified and correctly classified sequences. This analysis identified specific OGs associated with misclassification, providing insights into the evolutionary and functional characteristics that may contribute to classification errors. Statistical comparisons, such as the chi-square test, were employed to evaluate differences in COG distributions, highlighting significant patterns and associations.

#### 2.5.3. Motif Analysis: MEME Suite

To identify prevalent motifs within misclassified sequences, we used the MEME Suite [51], focusing on the Zero or One Occurrence per Sequence (ZOOPS) model. This analysis involved extracting motif information from text files and comparing motif occurrences between misclassified and correctly classified sequences. The tasks analyzed included solubility, localization, ion channels, transporters, and membrane proteins (MP). For each task, motifs were extracted from both misclassified and correctly classified sequences, and their occurrences were recorded. This allowed us to count the total number of sequences and motifs.

## 3. Results

The objective of this section is to evaluate the effectiveness of integrating secondary structure information into a pretrained Protein Language Model (PLM). We aim to determine if this integration enhances the model’s performance across various protein-related tasks. The integration utilized an encoder-decoder architecture based on the Ankh model, where the primary amino acid sequence is input to the model, and the model generates the corresponding secondary structure. This method implicitly incorporates secondary structure knowledge into the pretrained model, which has been trained on primary sequence data.

The performance of the resulting model, TooT-PLM-P2S, was assessed across multiple tasks using diverse datasets. Additionally, we conducted a detailed analysis of failure cases to understand the reasons for errors, employing three bioinformatics-based methods: Multiple Sequence Alignment (T-Coffee), Orthologous Groups (eggNOG) Analysis, and Motif Alignment and Search Tool (MEME Suite). This analysis provides insights into the model’s performance and highlights areas for improvement.

### 3.1. Overview of Results

In this section, we present an overview of our results, focusing on the comparative performance of the TooT-PLM-P2S model and the Ankh model. This summary establishes the foundation for detailed analyses in the subsequent sections.

We start by summarizing the cross-validation outcomes, emphasizing the models’ robustness and consistency across different folds. Then, we discuss the results from the independent test set to validate the model’s generalizability and real-world applicability. Further sections will provide an in-depth performance analysis categorized by secondary structure prediction (SSP) and non-SSP tasks, followed by a task-specific performance evaluation across seven distinct tasks.

#### 3.1.1. Cross-Validation Results

The results presented in Table 2 and Figure 4 compare the Ankh model and the TooT-PLM-P2S model using cross-validation. The Ankh model generally outperforms the TooT-PLM-P2S model across most tasks. However, there are notable exceptions where TooT-PLM-P2S demonstrates competitive or superior performance, particularly in tasks that may benefit more from the integration of secondary structure information. These findings highlight the strengths and limitations of each model in different protein-related tasks.

**Table 1.** Summary of Downstream Tasks for TooT-PLM-P2S Evaluation. The table summarizes the downstream tasks used to evaluate the TooT-PLM-P2S model. Each row details a specific task, including methodology, data source, number of protein sequences, and dataset splits for training, validation, and testing. Non-SSP tasks include Fluorescence Prediction (FluP), Solubility Prediction (SolP), Sub-cellular Localization Prediction (LocP), Ion Channel Prediction (IonP), Transporter Prediction (TranP), and Membrane Protein Prediction (MemP). SSP tasks involve Secondary Structure Prediction for three states (SSP3) and eight states (SSP8), reflecting the model’s performance in these areas.

| Task | Methodology | Data Source | #Proteins | #Classes | #Prot. Per Class(Pos./Neg.) | #Train/Valid./Test |
| --- | --- | --- | --- | --- | --- | --- |
| Non-SSP Tasks |  |  |  |  |  |  |
| FluP | Regression | Sarkisyan et al. [31] | 54,025 | - | - | 21,446/5,362/27,217 |
| SolP | Classification | DeepSol [32] | 71,421 | 2 | 29,972/41,449 | 62,478/6,942/1,999 |
| LocP | Classification | DeepLoc [33] | 13,961 | 10 | - | 8,945/2,248/2,768 |
| IonP | Classification | DeepIon [34] | 4,564 | 2 | 301/4,263 | 3,289/365/910 |
| TranP | Classification | TrSSP [35] | 1,560 | 2 | 900/660 | 1,242/138/180 |
| MemP | Classification | TooT-M [36] | 17,892 | 2 | 8,825/9,064 | 14,492/1,610/1,790 |
| SSP Tasks |  |  |  |  |  |  |
| SSP3 | Classification | NetSurfP-2.0 [28] | 11,361 | 3 | - | 8,678/2,170/513 |
| SSP8 | Classification | NetSurfP-2.0 [28] | 11,361 | 8 | - | 8,678/2,170/513 |

**Table 2.**
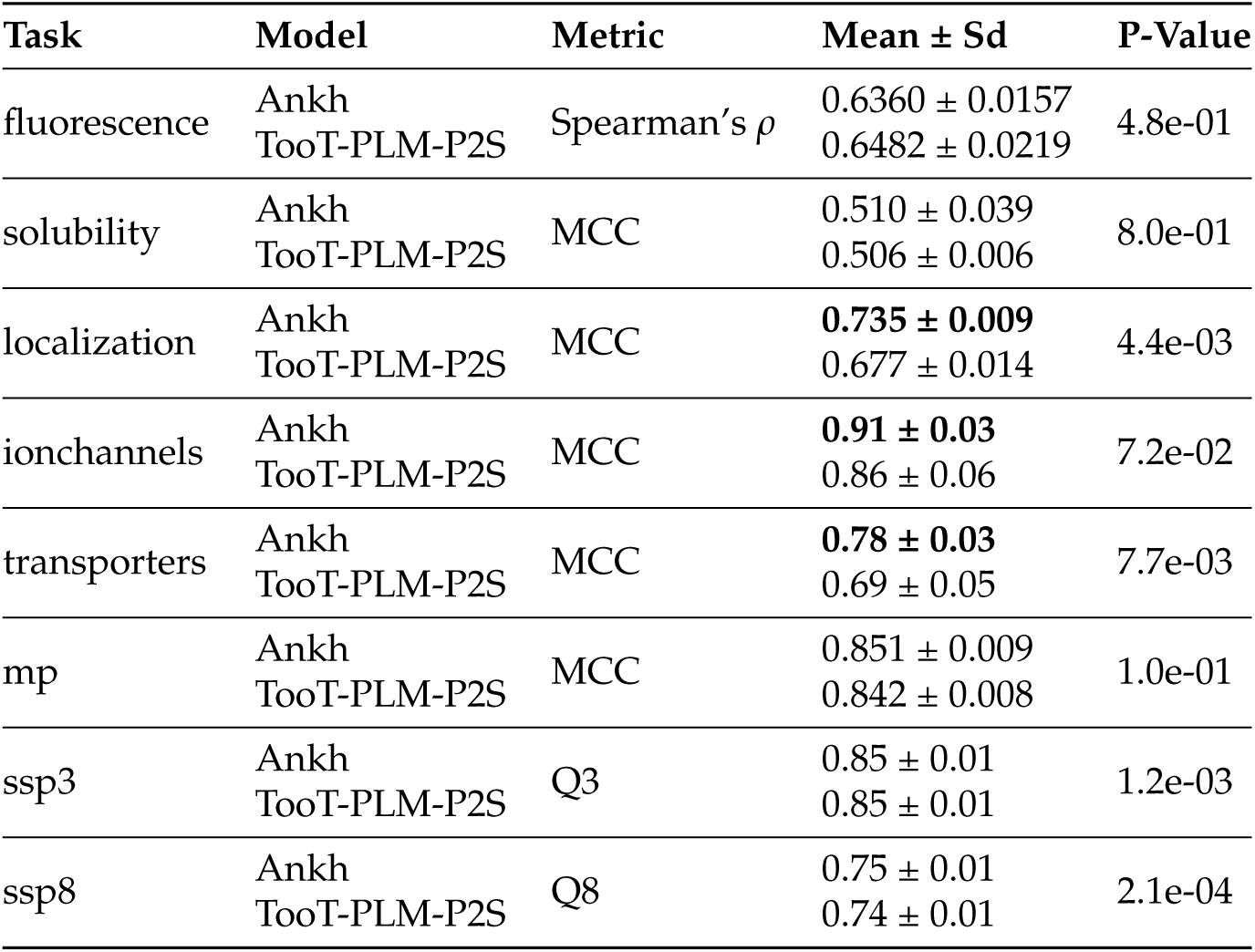
Comparative Performance Overview of Ankh and TooT-PLM-P2S on Cross-Validation.

**Figure 4.**
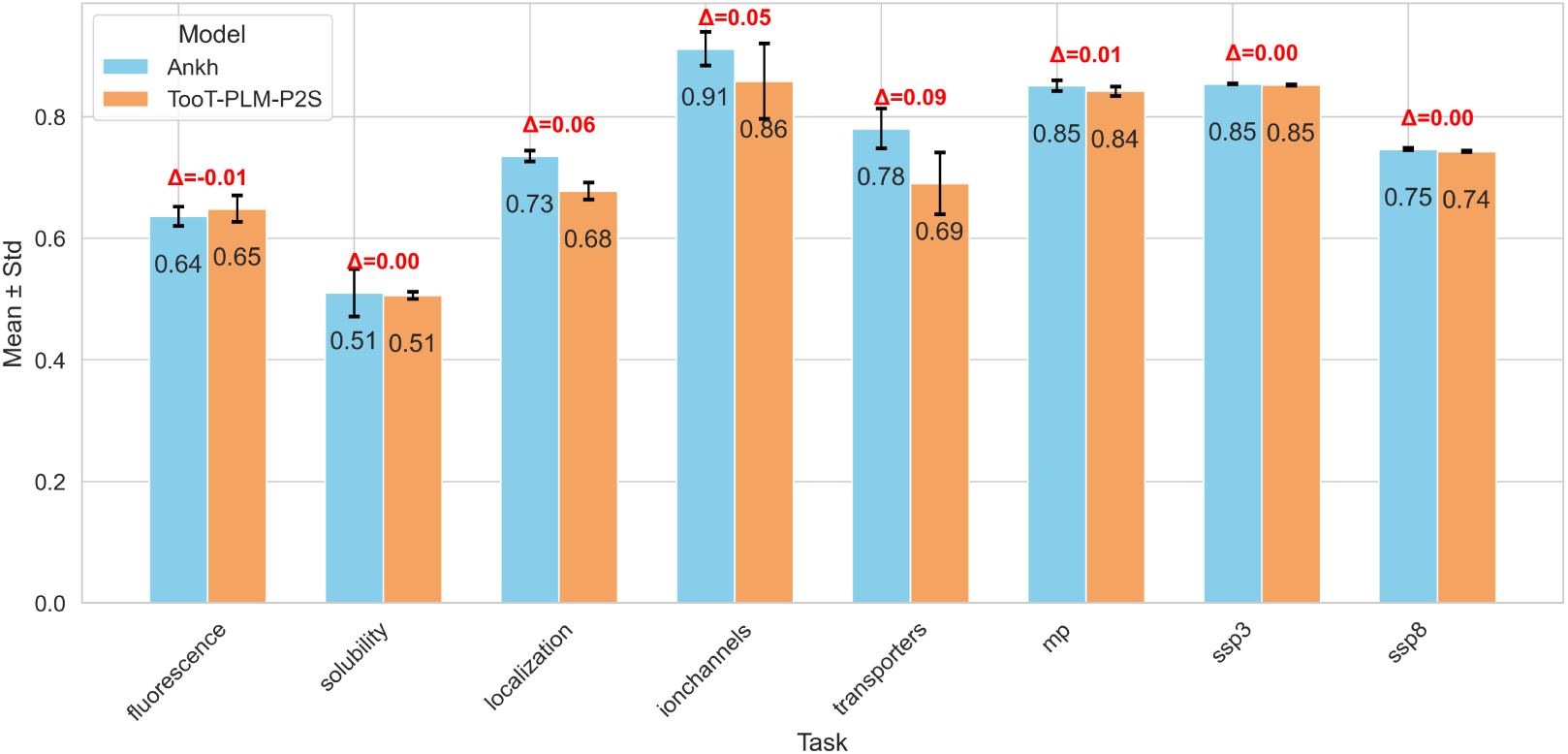
Comparative Performance Overview of Ankh and TooT-PLM-P2S on Cross-Validation This figure delineates a side-by-side comparative analysis of the Ankh and TooT-PLM-P2S models across the spectrum of prediction tasks. The x-axis enumerates the distinct tasks evaluated, while the y-axis quantifies the performance metrics, represented as mean values augmented with their respective standard deviations. Each task features a pair of bars, distinguished by colors as denoted in the figure’s legend, corresponding to the Ankh and TooT-PLM-P2S models. The bars reflect the average metric derived from a 5-fold cross-validation scheme, and the accompanying error bars delineate the standard deviation across the folds. The discrepancy observed between each pair of bars graphically portrays the performance differential between the two computational models.

This table provides an overview of the comparative performance between the Ankh and TooT-PLM-P2S models across various prediction tasks based on cross-validation datasets. For the Fluorescence Intensity Prediction Task, Spearman’s Correlation Coefficient (*ρ*) is used as the performance metric. The Matthews Correlation Coefficient (MCC) is employed for protein function prediction tasks, while Q3 and Q8 accuracy metrics are utilized for secondary structure prediction (SSP) tasks, indicating the proportion of correctly predicted secondary structure states. Notably, values in boldface within the table signify the better performance differences between the models. The p-value shows the statistical significance of the comparison between the two models. The threshold for statistical significance is set at a p-value of 0.05, meaning that p-values less than 0.05 are considered statistically significant.

In the fluorescence task, which is a regression task, the TooT-PLM-P2S model outperformed the Ankh model on the Spearman’s *ρ* metric, with a p-value of 0.48, indicating no statistically significant difference. For the solubility task, evaluated using the MCC, both models showed comparable performance, with a p-value of 0.80, also not statistically significant.

For the localization task, the Ankh model performed better than the TooT-PLM-P2S model, with a statistically significant p-value of 0.004. In the ion channels prediction task, the Ankh model showed superior performance with a p-value of 0.072, close to the significance threshold. For the transporters prediction task, the Ankh model outperformed the TooT-PLM-P2S model with a statistically significant p-value of 0.0077.

In the membrane proteins prediction task, the Ankh model showed better performance with a p-value of 0.10, not statistically significant. In the secondary structure prediction tasks, both three-state (SSP3) and eight-state (SSP8) tasks showed comparable results between the models. For SSP3, the p-value was 0.0012, and for SSP8, it was 0.00021, both indicating statistically significant differences.

In summary, while the Ankh model generally performs better across various tasks, the TooT-PLM-P2S model shows promise, particularly in fluorescence prediction. The statistical analysis, as indicated by the p-values, highlights significant and non-significant differences between the models.

#### 3.1.2. Independent Test Set Results

According to Table 3 and Table 5, the comparison between the Ankh model and the TooT-PLM-P2S model was evaluated on an independent test set across seven tasks. These tasks included secondary structure prediction (SSP) tasks assessed using Q3 and Q8 accuracy metrics. For classification tasks, the Matthews correlation coefficient (MCC) was used as the evaluation metric. For the fluorescence regression task, Spearman’s *ρ* was employed. The analysis focused on determining the performance differences between the models in various predictive tasks.

**Table 3.** Comparative performance of Ankh and TooT-PLM-P2S on the independent test set. This table presents a comparison of the performance of the Ankh model and the TooT-PLM-P2S model across various prediction tasks using an independent test set. For the fluorescence intensity prediction task, Spearman’s correlation coefficient (*ρ*) is used as the performance metric. Matthews correlation coefficient (MCC) is employed for protein function prediction tasks. Q3 and Q8 accuracy metrics are utilized for secondary structure prediction (SSP) tasks, indicating the proportion of correctly predicted secondary structure states. Values in boldface within the table indicate superior performance between the models.

| Task | Model | Metric | Value |
| --- | --- | --- | --- |
| fluorescence | TooT-PLM-P2S | Spearman’s $\rho$ | **0.6257** |
|  | Ankh |  | 0.6064 |
| solubility | TooT-PLM-P2S | MCC | 0.494 |
|  | Ankh |  | **0.517** |
| localization | TooT-PLM-P2S | MCC | 0.661 |
|  | Ankh |  | **0.766** |
| ionchannels | TooT-PLM-P2S | MCC | 0.82 |
|  | Ankh |  | **0.83** |
| transporters | TooT-PLM-P2S | MCC | 0.63 |
|  | Ankh |  | **0.82** |
| mp | TooT-PLM-P2S | MCC | **0.865** |
|  | Ankh |  | 0.845 |
| ssp3 | TooT-PLM-P2S | Q3 | 0.81 |
|  | Ankh |  | 0.81 |
| ssp8 | TooT-PLM-P2S | Q8 | 0.68 |
|  | Ankh |  | 0.68 |

The results on the independent test set align with those from the cross-validation analysis. Specifically, in the fluorescence task, the TooT-PLM-P2S model performed better on the Spearman’s *ρ* metric. However, for solubility, localization, ion channels, transporters, and membrane proteins tasks, the Ankh model demonstrated superior performance.

In the SSP3 task, both models showed comparable accuracy, while in the SSP8 task, the Ankh model had a slight advantage. For the ion channels task, the Ankh model also outperformed the TooT-PLM-P2S model slightly.

Overall, the independent test set results confirm the cross-validation findings, with the Ankh model generally excelling across most tasks, while the TooT-PLM-P2S model shows potential in the fluorescence task. Detailed analysis of non-SSP and SSP tasks follows in subsequent sections for a clearer understanding of model performance.

### 3.2. Performance By Category

In this section, we categorize performance results into non-SSP (non-secondary structure prediction) tasks and SSP (secondary structure prediction) tasks. This organization allows for a detailed and structured presentation of the results.

#### 3.2.1. Non-SSP Tasks

Non-SSP tasks include classification tasks such as solubility, localization, ion channels, transporters, and membrane proteins. We evaluated the Ankh and TooT-PLM-P2S models using several metrics: MCC, F1 score, recall, precision, and accuracy. We computed the average performance for each metric across all non-SSP tasks to facilitate a clear comparison of the overall effectiveness of the two models on these classification tasks. Table 4 compares the performance of the Ankh model and the TooT-PLM-P2S model across these various tasks.

**Table 4.** Non-SSP tasks overview of the TooT-PLM-P2S and Ankh model comparison. This table presents a comparison of non-SSP tasks between the Ankh and TooT-PLM-P2S models on both the cross-validation and test sets. The p-values indicate the statistical significance of the results. The threshold for statistical significance is set at a p-value of 0.05, meaning that p-values less than 0.05 are considered statistically significant. MCC stands for Matthews correlation coefficient.

| Metric | Model | Cross-Validation | Test Set | P-Value |
| --- | --- | --- | --- | --- |
| MCC | Ankh | $0.757 \pm 0.138$ | 0.756 | 0.0566 |
| | TooT-PLM-P2S | $0.714 \pm 0.128$ | 0.693 | |
| F1 | Ankh | $0.844 \pm 0.081$ | 0.838 | 0.0165 |
| | TooT-PLM-P2S | $0.807 \pm 0.083$ | 0.799 | |
| Recall | Ankh | $0.862 \pm 0.069$ | 0.821 | 0.0392 |
| | TooT-PLM-P2S | $0.791 \pm 0.102$ | 0.812 | |
| Precision | Ankh | $0.837 \pm 0.092$ | 0.866 | 0.9780 |
| | TooT-PLM-P2S | $0.838 \pm 0.093$ | 0.797 | |
| Accuracy | Ankh | $0.867 \pm 0.090$ | 0.878 | 0.2482 |
| | TooT-PLM-P2S | $0.850 \pm 0.093$ | 0.841 | |

Overall, the Ankh model outperformed the TooT-PLM-P2S model on several metrics during cross-validation, specifically MCC, F1 score, recall, and accuracy. However, for the precision metric, both models demonstrated comparable performance. These cross-validation results indicate that the Ankh model generally provides superior performance compared to the TooT-PLM-P2S model, except in precision.

The p-values associated with these results show statistical significance for the F1 score and recall metrics, as they are less than 0.05. However, for MCC, precision, and accuracy, the p-values are not less than 0.05, indicating that the differences in these metrics are not statistically significant.

Additionally, the performance of the two models was compared on the test set for the non-SSP tasks. Consistent with the cross-validation results, the Ankh model demonstrated better performance than the TooT-PLM-P2S model on the test set.

In summary, the Ankh model generally exhibits superior performance over the TooT-PLM-P2S model in both cross-validation and test set evaluations, except for precision during cross-validation where both models performed comparably. The statistical significance of the results is confirmed for the F1 score and recall metrics.

#### 3.2.2. SSP Tasks

In this section, we compare the performance of the Ankh model and the TooT-PLM-P2S model on SSP tasks, specifically SSP3 and SSP8. Table 5 presents the evaluation metrics, including F1 score, recall, precision, and accuracy, computed as averages across cross-validation and test sets. The p-values are also included to assess the statistical significance of the results.

**Table 5.**
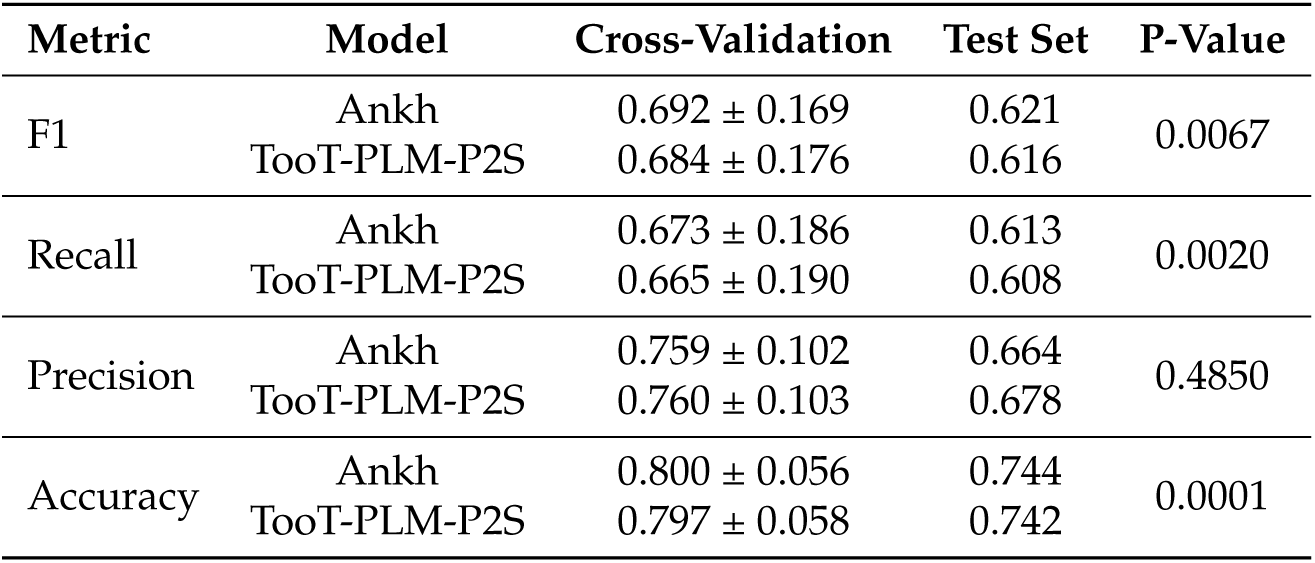
SSP tasks overview of the TooT-PLM-P2S and Ankh model comparison. This table presents a comparison of SSP tasks between the Ankh and TooT-PLM-P2S models, displaying averages of F1 score, recall, precision, and accuracy for both SSP3 and SSP8 across cross-validation and test sets. The p-values indicate the statistical significance of these results. The threshold for statistical significance is set at a p-value of 0.05, meaning that p-values less than 0.05 are considered statistically significant.

| Metric | Model | Cross-Validation | Test Set | P-Value |
| --- | --- | --- | --- | --- |
| F1 | Ankh | $0.692 \pm 0.169$ | 0.621 | 0.0067 |
| | TooT-PLM-P2S | $0.684 \pm 0.176$ | 0.616 | |
| Recall | Ankh | $0.673 \pm 0.186$ | 0.613 | 0.0020 |
| | TooT-PLM-P2S | $0.665 \pm 0.190$ | 0.608 | |
| Precision | Ankh | $0.759 \pm 0.102$ | 0.664 | 0.4850 |
| | TooT-PLM-P2S | $0.760 \pm 0.103$ | 0.678 | |
| Accuracy | Ankh | $0.800 \pm 0.056$ | 0.744 | 0.0001 |
| | TooT-PLM-P2S | $0.797 \pm 0.058$ | 0.742 | |

Based on the results in Table 5, the Ankh model outperformed the TooT-PLM-P2S model in terms of F1 score and recall. The TooT-PLM-P2S model demonstrated better performance in precision. Both models showed comparable results in accuracy.

The statistical significance analysis indicated by p-values reveals significant differences in F1 score, recall, and accuracy between the models, with p-values less than 0.05. For precision, where the TooT-PLM-P2S model outperformed the Ankh model, the p-value is greater than 0.05, indicating the result is not statistically significant.

Overall, these findings suggest that while the Ankh model excels in F1 score and recall for SSP tasks, both models perform similarly in terms of accuracy. The TooT-PLM-P2S model shows an edge in precision, although this result is not statistically significant.

### 3.3. Task-Specific Results

This section provides detailed observations from seven different tasks: fluorescence prediction, solubility, subcellular localization, ion channel classification, transporter classification, membrane protein classification, and secondary structure prediction. Examining each task individually offers a comprehensive understanding of the performance differences between the Ankh and TooT-PLM-P2S models. This analysis highlights specific strengths and weaknesses of each model across various biological prediction tasks.

#### 3.3.1. Fluorescence Prediction

We compared the performance of the Ankh model and the TooT-PLM-P2S model for fluorescence prediction using Spearman’s correlation coefficient (*ρ*). The TooT-PLM-P2S model showed better performance in terms of mean, standard deviation, maximum, and median *ρ* values. The mean *ρ* for TooT-PLM-P2S was higher, and the standard deviation was also greater. The maximum *ρ* achieved by TooT-PLM-P2S was higher, and the median *ρ* was better for TooT-PLM-P2S. However, the differences in performance between the two models were not statistically significant, with a p-value of 0.48. The detailed comparison metrics are presented in Table 6.

**Table 6.** Fluorescence prediction comparison between TooT-PLM-P2S and Ankh models. This table presents the comparison of fluorescence prediction performance between the TooT-PLM-P2S and Ankh models, using Spearman’s correlation coefficient (*ρ*) on cross-validation. The table includes mean, standard deviation, maximum, minimum, and median values from the cross-validation results, along with the p-value indicating the statistical significance of the comparison. The threshold for statistical significance is set at a p-value of 0.05, meaning that p-values less than 0.05 are considered statistically significant. Boldface values denote higher performance in the comparison between the two models.

| Model | Mean ± Sd | Max | Min | Median | P-Value |
| --- | --- | --- | --- | --- | --- |
| Ankh | 0.6360 ± 0.0157 | 0.6533 | 0.6160 | 0.6375 | 4.8e-01 |
| TooT-PLM-P2S | 0.6482 ± 0.0219 | 0.6742 | 0.6139 | 0.6523 | |

#### 3.3.2. Solubility Prediction

We compared the Ankh and TooT-PLM-P2S models for solubility prediction using accuracy, precision, recall, F1 score, and Matthews correlation coefficient (MCC). The results, shown in Table 7, highlight the performance metrics for each model in this task.

**Table 7.** Solubility Prediction: Performance Comparison between the Ankh and TooT-PLM-P2S models. This table presents the performance comparison of the Ankh and TooT-PLM-P2S models for solubility prediction, using cross-validation. The metrics reported include accuracy, precision, recall, F1 score, and Matthews correlation coefficient (MCC). Each metric’s p-value is provided to indicate statistical significance. The threshold for statistical significance is set at a p-value of 0.05, meaning that p-values less than 0.05 are considered statistically significant. Boldface values denote the higher performance between the models.

|  | Accuracy | Precision | Recall | F1 | MCC |
| --- | --- | --- | --- | --- | --- |
| Ankh | 0.745 ± 0.048 | 0.687 ± 0.088 | 0.779 ± 0.117 | **0.719 ± 0.010** | 0.510 ± 0.039 |
| TooT-PLM-P2S | 0.762 ± 0.003 | 0.754 ± 0.019 | 0.641 ± 0.037 | 0.692 ± 0.014 | 0.506 ± 0.006 |
| P-Value | 4.50e-01 | 2.30e-01 | 1.10e-01 | 4.60e-03 | 8.00e-01 |

The TooT-PLM-P2S model achieved higher accuracy and precision than the Ankh model. However, the Ankh model showed higher recall and F1 score. Both models had comparable MCC values. The p-values indicate that the differences in accuracy, precision, recall, and MCC are not statistically significant (p-value > 0.05). Only the difference in F1 score is statistically significant (p-value < 0.05).

3.3.3. Sub-cellular Localization Prediction

In this subsection, we compare the performance of the Ankh model and the TooT-PLM-P2S model for subcellular localization prediction. Table 8 summarizes the evaluation metrics, including accuracy, precision, recall, F1 score, and MCC. Each metric is accompanied by its respective p-value to assess statistical significance.

**Table 8.** Subcellular Localization Prediction: Performance Comparison between the Ankh and TooT-PLM-P2S models. This table presents the performance comparison of the Ankh and TooT-PLM-P2S models for subcellular localization prediction, using cross-validation. The metrics reported include accuracy, precision, recall, F1 score, and Matthews correlation coefficient (MCC). Each metric’s p-value is provided to indicate statistical significance. The threshold for statistical significance is set at a p-value of 0.05, meaning that p-values less than 0.05 are considered statistically significant. Boldface values denote the higher performance between the models.

|  | Accuracy | Precision | Recall | F1 | MCC |
| --- | --- | --- | --- | --- | --- |
| Ankh | **0.786 ± 0.007** | 0.792 ± 0.012 | **0.786 ± 0.007** | **0.781 ± 0.009** | **0.735 ± 0.009** |
| TooT-PLM-P2S | 0.739 ± 0.010 | 0.753 ± 0.022 | 0.739 ± 0.010 | 0.735 ± 0.009 | 0.677 ± 0.014 |
| P-Value | 2.90e-03 | 5.70e-02 | 2.90e-03 | 3.70e-03 | 4.40e-03 |

According to Table 8, the Ankh model outperformed the TooT-PLM-P2S model across all evaluation metrics. The p-values for each metric are less than 0.05, indicating statistically significant differences in performance between the two models.

#### 3.3.4. Ion Channel Classification

This section compares the performance of the Ankh and TooT-PLM-P2S models for ion channel classification. The evaluation metrics, including accuracy, precision, recall, F1 score, and Matthews correlation coefficient (MCC), are summarized in Table 9.

**Table 9.** Ion Channel Classification: Performance Comparison between the Ankh and TooT-PLM-P2S models. This table presents the performance comparison of the Ankh and TooT-PLM-P2S models for ion channel classification, using cross-validation. The metrics reported include accuracy, precision, recall, F1 score, and Matthews correlation coefficient (MCC). Each metric’s p-value is provided to indicate statistical significance. The threshold for statistical significance is set at a p-value of 0.05, meaning that p-values less than 0.05 are considered statistically significant. Boldface values denote the higher performance between the models.

|  | Accuracy | Precision | Recall | F1 | MCC |
| --- | --- | --- | --- | --- | --- |
| Ankh | 0.99 ± 0.003 | 0.95 ± 0.037 | **0.88 ± 0.035** | 0.92 ± 0.026 | 0.91 ± 0.028 |
| TooT-PLM-P2S | 0.98 ± 0.007 | 0.98 ± 0.023 | 0.77 ± 0.098 | 0.86 ± 0.064 | 0.86 ± 0.062 |
| P-Value | 7.7e-02 | 3.0e-01 | 2.0e-02 | 6.0e-02 | 7.2e-02 |

The Ankh and TooT-PLM-P2S models show comparable accuracy, with no statistically significant difference. The TooT-PLM-P2S model achieves higher precision, but this difference is also not statistically significant. The Ankh model has higher recall, though the difference is not statistically significant. The Ankh model outperforms the TooT-PLM-P2S model in F1 score and MCC, with both differences being statistically significant.

#### 3.3.5. Transporter Classification

This section compares the performance of the Ankh and TooT-PLM-P2S models for transporter classification. The evaluation metrics, including accuracy, precision, recall, F1 score, and MCC, are summarized in Table 10.

**Table 10.** Transporter Classification: Performance Comparison between the Ankh and TooT-PLM-P2S models. This table presents the performance comparison of the Ankh and TooT-PLM-P2S models for transporter classification, using cross-validation. The metrics reported include accuracy, precision, recall, F1 score, and Matthews correlation coefficient (MCC). Each metric’s p-value is provided to indicate statistical significance. The threshold for statistical significance is set at a p-value of 0.05, meaning that p-values less than 0.05 are considered statistically significant. Boldface values denote the higher performance between the models.

|  | Accuracy | Precision | Recall | F1 | MCC |
| --- | --- | --- | --- | --- | --- |
| Ankh | **0.89 ± 0.02** | **0.86 ± 0.03** | 0.90 ± 0.01 | **0.88 ± 0.02** | **0.78 ± 0.03** |
| TooT-PLM-P2S | 0.84 ± 0.03 | 0.79 ± 0.05 | 0.88 ± 0.03 | 0.83 ± 0.03 | 0.69 ± 0.05 |
| P-Value | 1.1e-02 | 2.6e-02 | 8.1e-02 | 5.8e-03 | 7.7e-03 |

The Ankh model demonstrates higher accuracy than the TooT-PLM-P2S model, but this difference is not statistically significant. Precision is also higher for the Ankh model, yet this difference is not statistically significant either. However, the Ankh model significantly outperforms the TooT-PLM-P2S model in recall, F1 score, and MCC, as indicated by the statistically significant p-values.

#### 3.3.6. Membrane Protein Classification

In this section, we compare the performance of the Ankh and TooT-PLM-P2S models for membrane protein classification. The evaluation metrics, including accuracy, precision, recall, F1 score, and MCC, along with their respective p-values to assess statistical significance, are presented in Table 11.

**Table 11.**
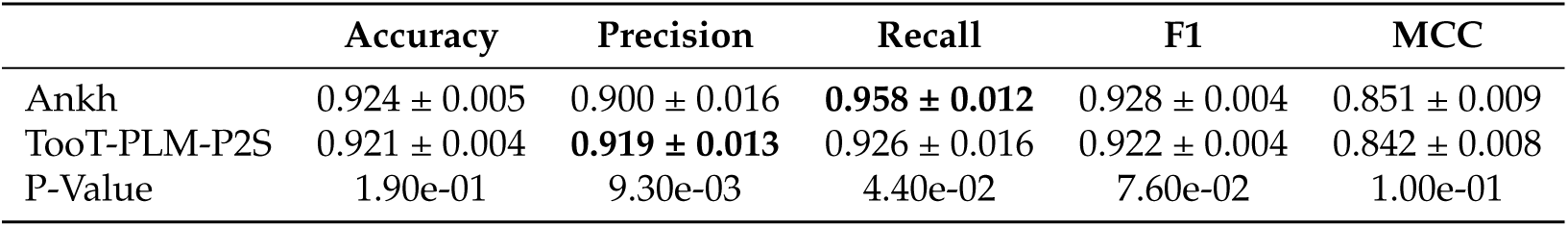
Membrane Protein Classification: Performance Comparison between the Ankh and TooT-PLM-P2S model. This table presents the performance comparison of the Ankh and TooT-PLM-P2S models for membrane protein classification. The evaluation metrics include accuracy, precision, recall, F1 score, and MCC, with corresponding p-values indicating statistical significance. The threshold for statistical significance is set at a p-value of 0.05, meaning that p-values less than 0.05 are considered statistically significant. Boldface values highlight the higher performance between the models.

|  | Accuracy | Precision | Recall | F1 | MCC |
| --- | --- | --- | --- | --- | --- |
| Ankh | 0.924 ± 0.005 | 0.900 ± 0.016 | **0.958 ± 0.012** | 0.928 ± 0.004 | 0.851 ± 0.009 |
| TooT-PLM-P2S | 0.921 ± 0.004 | **0.919 ± 0.013** | 0.926 ± 0.016 | 0.922 ± 0.004 | 0.842 ± 0.008 |
| P-Value | 1.90e-01 | 9.30e-03 | 4.40e-02 | 7.60e-02 | 1.00e-01 |

The accuracy of the Ankh and TooT-PLM-P2S models is comparable, with a p-value greater than 0.05, indicating no statistically significant difference. The F1 scores of the two models are also comparable, but the p-value for the F1 score is less than 0.05, suggesting statistical significance. The Ankh model shows better performance in recall and MCC compared to the TooT-PLM-P2S model, although the p-values for these metrics are greater than 0.05, indicating that the differences are not statistically significant. The TooT-PLM-P2S model outperforms the Ankh model in precision, with a p-value less than 0.05, indicating statistical significance.

#### 3.3.7. Secondary Structure Prediction

We analyzed the performance of the Ankh and TooT-PLM-P2S models across various non-SSP tasks and now evaluate their performance on Secondary Structure Prediction (SSP) tasks. This section compares the models’ ability to predict secondary structure elements of proteins, focusing on SSP-3 and SSP-8 prediction tasks.

SSP-3 prediction involves categorizing protein residues into three secondary structure classes: alpha-helix, beta-strand, and coil. SSP-8 prediction entails a more granular classification into eight distinct secondary structure states.

We present the results for both SSP-3 and SSP-8 predictions, showcasing the evaluation metrics including F1 score, recall, precision, and accuracy for each model. These metrics provide insights into the models’ performance in predicting secondary structures, a critical task in bioinformatics.

##### SSP3 Prediction

We evaluate the performance of the Ankh and TooT-PLM-P2S models on SSP-3 prediction tasks. Table 12 provides a comparison of the models using four evaluation metrics: accuracy, precision, recall, and F1 score. Each metric is accompanied by its respective p-value to assess statistical significance.

**Table 12.** SSP-3 Prediction: Performance Comparison between the Ankh and TooT-PLM-P2S models. This table presents the performance comparison of the Ankh and TooT-PLM-P2S models for SSP-3 prediction. The evaluation metrics include accuracy, precision, recall, and F1 score, each with their respective p-values to indicate statistical significance. The threshold for statistical significance is set at a p-value of 0.05, meaning that p-values less than 0.05 are considered statistically significant.

|  | Accuracy | Precision | Recall | F1 |
| --- | --- | --- | --- | --- |
| Ankh | 0.83 ± 0.04 | 0.83 ± 0.05 | 0.82 ± 0.04 | 0.83 ± 0.05 |
| TooT-PLM-P2S | 0.83 ± 0.04 | 0.83 ± 0.05 | 0.82 ± 0.04 | 0.82 ± 0.05 |
| P-Value | 8.9e-01 | 7.2e-01 | 2.7e-01 | 6.0e-01 |

According to Table 12, there is no discernible difference between the Ankh and TooT-PLM-P2S models across all evaluation metrics. Both models consistently achieve a value of 82% for accuracy, precision, recall, and F1 score. The p-values for each metric are greater than 0.05, indicating that these differences are not statistically significant. This demonstrates that the performance of the Ankh and TooT-PLM-P2S models is equivalent on SSP-3 prediction tasks, with no significant variation observed in any of the metrics.

##### SSP8 Prediction

We evaluate the performance of the Ankh and TooT-PLM-P2S models on SSP-8 prediction tasks. Table 13 provides a comparison of the models using four evaluation metrics: accuracy, precision, recall, and F1 score, with corresponding p-values to assess statistical significance.

**Table 13.** SSP-8 Prediction: Performance Comparison between the Ankh and TooT-PLM-P2S models. This table presents the performance comparison of the Ankh and TooT-PLM-P2S models for SSP-8 prediction. The evaluation metrics include accuracy, precision, recall, and F1 score, each with their respective p-values to indicate statistical significance. The threshold for statistical significance is set at a p-value of 0.05, meaning that p-values less than 0.05 are considered statistically significant. Boldface values highlight the higher performance between the models.

|  | Accuracy | Precision | Recall | F1 |
| --- | --- | --- | --- | --- |
| Ankh | **0.71 ± 0.05** | 0.59 ± 0.11 | **0.46 ± 0.06** | **0.49 ± 0.07** |
| TooT-PLM-P2S | 0.71 ± 0.05 | 0.61 ± 0.10 | 0.45 ± 0.05 | 0.48 ± 0.07 |
| P-Value | 1.8e-05 | 2.3e-01 | 2.1e-05 | 1.0e-04 |

According to Table 13, the Ankh model outperforms the TooT-PLM-P2S model in all evaluation metrics except precision. Specifically, the Ankh model achieves higher values in accuracy, recall, and F1 score. The p-values for these metrics are less than 0.05, indicating that the differences are statistically significant. This demonstrates the superior performance of the Ankh model over the TooT-PLM-P2S model in terms of accuracy, recall, and F1 score for SSP-8 prediction.

Conversely, the TooT-PLM-P2S model outperforms the Ankh model in precision. However, the p-value for precision is greater than 0.05, indicating that this difference is not statistically significant. Therefore, while the Ankh model shows statistically significant improvements in accuracy, recall, and F1 score, the advantage of the TooT-PLM-P2S model in precision is not statistically significant.

## 4. Discussion

This section interprets and analyzes the key findings of our study, which evaluated the integration of secondary structure information into a pretrained protein language model. The results indicate that while the Ankh model generally outperformed the TooT-PLM-P2S model across various tasks, the integrated model demonstrated promise, particularly in fluorescence prediction and precision metrics.

We examine the performance differences between the models and their implications for protein-related tasks. This discussion includes an analysis of failure cases using bioinformatics tools such as multiple sequence alignment with T-Coffee, functional annotation with eggNOG, and motif analysis with MEME Suite.

### 4.1. Key Findings and Interpretations

This section presents the key findings from our study, which evaluated the integration of secondary structure information into a pre-trained protein language model (PLM) using the TooT-PLM-P2S model compared to the Ankh model. The results are interpreted for each primary task investigated.

For fluorescence prediction, the TooT-PLM-P2S model exhibited a higher mean Spearman’s correlation coefficient compared to the Ankh model. However, this improvement was not statistically significant, indicating that the observed advantage might be due to random variations rather than a genuine enhancement in model performance.

In solubility prediction, the Ankh model had a significantly higher F1 score compared to the TooT-PLM-P2S model. Other metrics such as accuracy, precision, and recall showed mixed results, but these differences were not statistically significant, suggesting that the observed variations could be due to random chance rather than a real difference in model performance.

For subcellular localization prediction, the Ankh model significantly outperformed the TooT-PLM-P2S model across almost all evaluation metrics, including MCC, F1, recall, and accuracy. This indicates that the integration of secondary structure information did not enhance performance for this task and may require further refinement.

In ion channel classification, only the recall metric showed statistically significant results, with the Ankh model outperforming the TooT-PLM-P2S model. While the Ankh model also demonstrated superior performance in F1 score and MCC, these differences were not statistically significant. The TooT-PLM-P2S model performed better in precision, but this was also not statistically significant, indicating that the incorporation of secondary structure information might improve specific metrics but not overall performance.

The Ankh model demonstrated better results across almost all metrics for transporter classification, including MCC, F1 score, recall, and accuracy. These differences were statistically significant, indicating that primary sequence information is more effective for this task compared to the integration of secondary structure information in the TooT-PLM-P2S model.

In membrane protein classification, the TooT-PLM-P2S model exhibited better performance in precision compared to the Ankh model, and this difference was statistically significant. Conversely, the Ankh model outperformed TooT-PLM-P2S in recall with significant results. The other metrics, such as MCC and accuracy, did not show statistically significant differences, indicating that secondary structure integration does not consistently improve performance across all metrics.

For secondary structure prediction (SSP3 and SSP8), all results for SSP3 were not statistically significant across all metrics. For SSP8 prediction, the Ankh model outperformed the TooT-PLM-P2S model with statistically significant differences in the F1 and recall metrics. The other metrics did not show significant differences.

These findings highlight the complexities of integrating secondary structure information into protein language models. The TooT-PLM-P2S model shows potential benefits, particularly in the precision of membrane protein classification, where it shows statistically significant improvement. However, further refinement and optimization are necessary to confirm and expand upon these benefits across other tasks and metrics.

### 4.2. Performance of TooT-PLM-P2S in Fluorescence Prediction

The TooT-PLM-P2S model did not show a statistically significant improvement in fluorescence prediction compared to the Ankh model. While the TooT-PLM-P2S model had a higher mean Spearman’s correlation coefficient, the lack of statistical significance indicates that these results could be due to random variations rather than a genuine enhancement from integrating secondary structure information. This suggests that while secondary structure data might provide additional contextual cues, further validation with larger datasets and more robust methods is needed to confirm its impact on predictive accuracy. Thus, the potential benefits of integrating secondary structure information remain inconclusive without statistically significant evidence.

### 4.3. Performance of TooT-PLM-P2S in Solubility Prediction

In solubility prediction, the only metric with a statistically significant result is the F1 score, where the Ankh model outperforms the TooT-PLM-P2S model. This indicates that integrating secondary structure information did not improve the overall predictive performance of the TooT-PLM-P2S model in this task. While the TooT-PLM-P2S model showed higher accuracy and precision, these differences were not statistically significant, suggesting that any observed advantages could be due to random variations rather than true enhancements in model performance. Therefore, the Ankh model’s superior F1 score highlights its better reliability in solubility prediction without the need for secondary structure integration.

### 4.4. Impact on Secondary Structure Prediction

The integration of secondary structure information into the PLM with the TooT-PLM-P2S model aimed to enhance predictive capabilities by providing additional structural context. However, the results indicated that this integration did not lead to superior performance in secondary structure prediction compared to the Ankh model. One potential reason for this outcome is that the current method of incorporating secondary structure data may not effectively leverage the complex relationships between primary and secondary structures. If the integration approach does not accurately model these relationships, it may fail to offer significant advantages over simpler, well-optimized models like Ankh.

Additionally, the dataset used for training TooT-PLM-P2S, sourced from the NetSurf project [28], may have been insufficient in terms of sequence variety and number. This limitation can hinder the model’s ability to effectively learn and generalize secondary structure information. The insufficient size of the dataset could have impeded the model’s capacity to grasp intricate patterns necessary for accurate secondary structure prediction. Future models might achieve better performance by leveraging more extensive and diverse datasets that provide a broader range of protein sequences. Expanding and refining training datasets is critical for developing robust predictive models in bioinformatics.

### 4.5. Enrichment Analysis of Misclassifications

Understanding misclassifications by machine learning models is crucial for enhancing their performance and reliability in protein classification. Misclassifications reveal model limitations and guide iterative improvements. This Enrichment Analysis (EA) of misclassified sequences by the Ankh and TooT-PLM-P2S models utilized T-Coffee, eggNOG, and the Motif Alignment and Search Tool (MAST) to investigate alignment patterns, evolutionary contexts, and motif characteristics. These findings provide a thorough examination of misclassifications, informing future model refinements. Table 14 presents the number of common correctly classified and misclassified sequences for both models analyzed in this section.

**Table 14.** Common Correctly Classified and Misclassified Sequences for Both Models. This table presents the number of common correctly classified and misclassified sequences for both the Ankh and TooT-PLM-P2S models across different tasks. The tasks include solubility, localization, ion channels, transporters, and membrane proteins (MP). The table shows the number of sequences misclassified and correctly classified in each task, along with the total number of sequences analyzed per task. The total row at the end summarizes the overall counts for all tasks combined.

| Task | Misclassified Sequences | Correctly Classified Sequences | Total Sequences |
| --- | --- | --- | --- |
| Solubility | 294 | 1293 | 1587 |
| Localization | 215 | 1206 | 1421 |
| Ionchannels | 13 | 888 | 901 |
| Transporters | 9 | 146 | 155 |
| MP | 93 | 1619 | 1712 |
| Total | 624 | 5152 | 5776 |

#### 4.5.1. T-Coffee Sequence Alignment

The T-Coffee sequence alignment analysis, shown in Figure 6, highlights significant variations in average sequence identities between misclassified and correctly classified sequences across different protein classification tasks. These variations illustrate the challenges faced by the models in accurately classifying certain sequences, providing critical insights into the sequence regions or motifs that may lead to misclassification. This information is essential for understanding and addressing specific issues that impact model performance.

**Figure 5.**
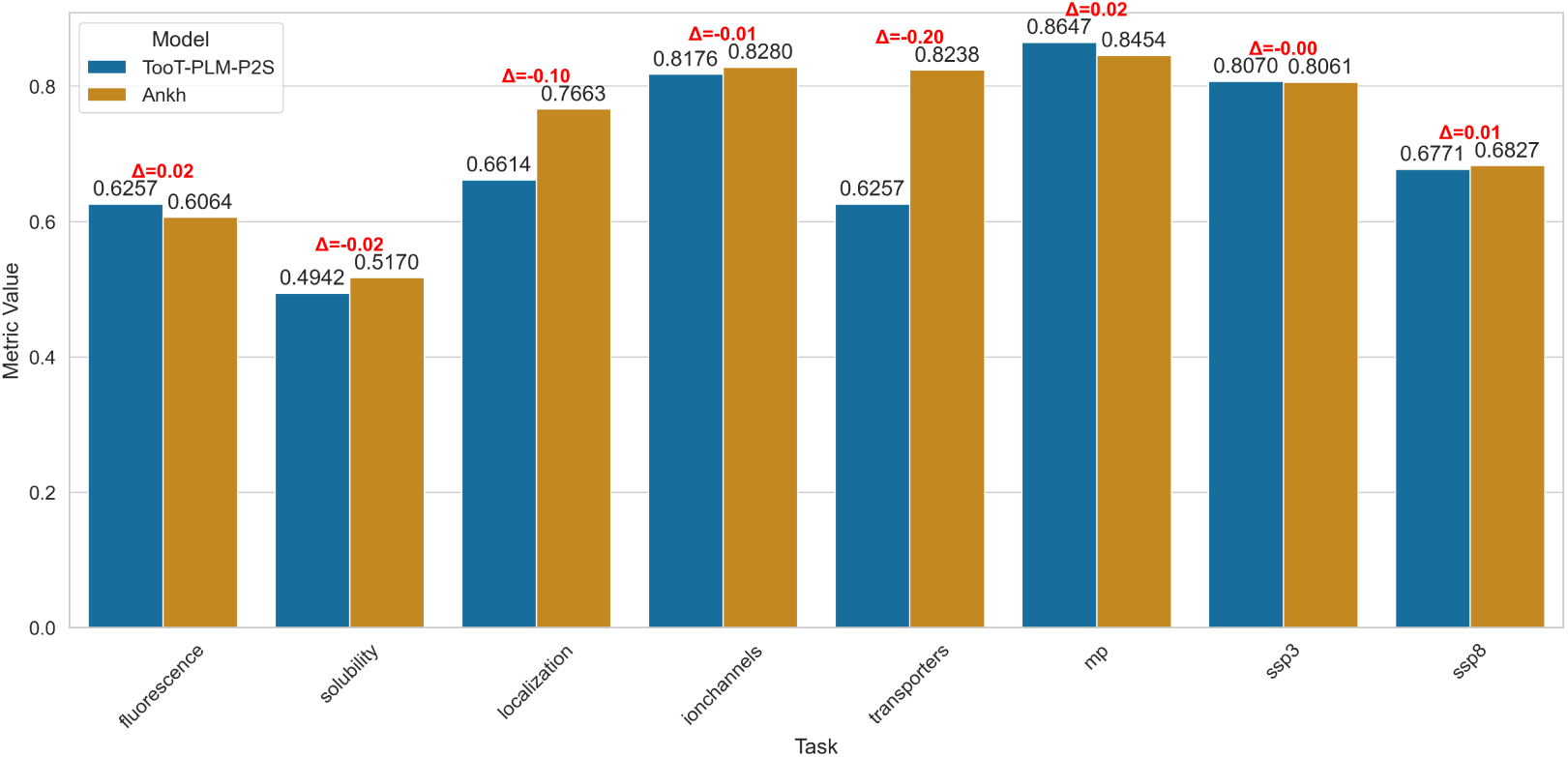
Comparative performance overview of Ankh and TooT-PLM-P2S on the test set. This figure provides a side-by-side comparative analysis of the Ankh model and the TooT-PLM-P2S model across a range of prediction tasks on the independent test set. The x-axis enumerates the distinct tasks evaluated, while the y-axis quantifies the performance metrics. Each task features a pair of bars, distinguished by colors as denoted in the figure’s legend, corresponding to the performance of the Ankh and TooT-PLM-P2S models. The visual representation highlights the performance differences, making it easier to identify which model performs better on each task.

**Figure 6.**
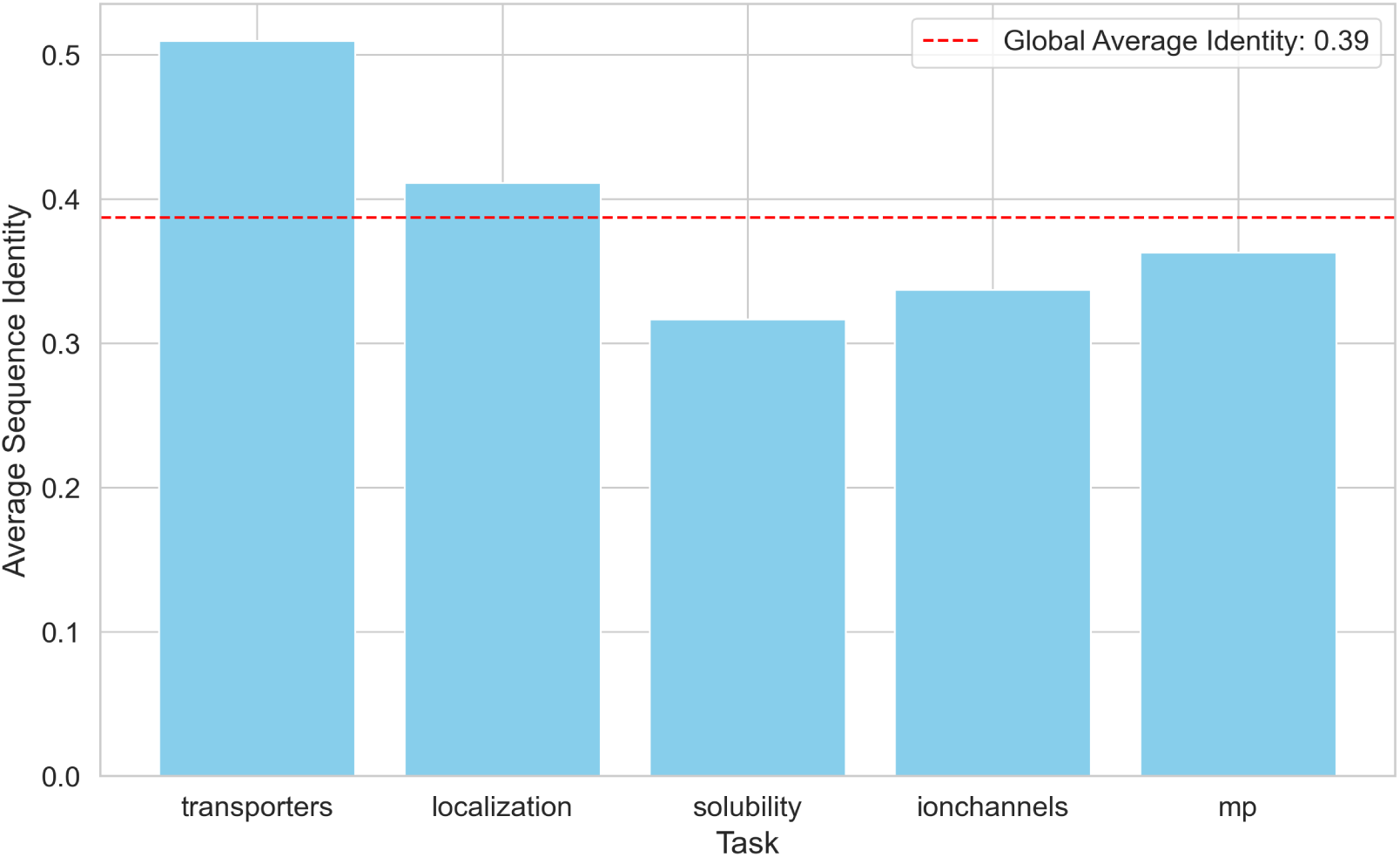
Average Sequence Identity for Misclassified vs Correctly Classified Sequences. This bar plot illustrates the average sequence identity between misclassified sequences and correctly classified sequences across five different protein classification tasks: transporters, localization, solubility, ion channels, and membrane proteins (MP). The overall global average identity is also presented, reflecting the general trend across all tasks.

For the transporters task, the average sequence identity is approximately 0.51, indicating substantial similarity between misclassified and correctly classified sequences. This suggests that misclassifications may arise from subtle differences not captured by current model features, possibly in critical protein regions affecting function. In the solubility task, the average identity is about 0.32, reflecting greater divergence and suggesting complex factors influencing solubility that are not apparent in primary sequences alone. Tasks such as localization and ion channels show intermediate identities (around 0.41 and 0.34), indicating moderate similarity and highlighting the models’ difficulty with weak or confounded functional signals. The overall average identity of 0.39 across all tasks underscores the need for models to address both conserved and variable sequence regions. Integrating complex features like secondary and tertiary structures, evolutionary conservation patterns, and biochemical properties could enhance model accuracy and robustness, as illustrated in Figure 6.

#### 4.5.2. Functional Annotation with eggNOG

The eggNOG analysis of misclassified sequences provides data on the distribution of correctly classified and misclassified sequences across various COG categories, as shown in Table 15. The ’S’ category, representing proteins with unknown functions, has the highest number of misclassified sequences (158), indicating that the model struggles with proteins lacking well-defined functions. Categories such as ’K’ (Transcription) and ’T’ (Signal transduction mechanisms) also suggest that the complexity and diversity within these groups confuse the models. These findings highlight areas for potential model improvement.

**Table 15.** EggNOG Analysis of Misclassified Sequences. This table presents the results of the eggNOG analysis, detailing the distribution of correctly classified and misclassified protein sequences across various Clusters of Orthologous Groups (COG) categories. Each row represents a specific COG category, providing insights into RNA processing, amino acid transport, carbohydrate metabolism, translation, transcription, replication, post-translational modification, inorganic ion transport, and signal transduction mechanisms.

| COG Category | Description | Correctly Classified | Misclassified | Percentage Misclassified |
| --- | --- | --- | --- | --- |
| A | RNA processing and modification | 130 | 21 | 13.89% |
| E | Amino acid transport and metabolism | 132 | 21 | 13.73% |
| G | Carbohydrate transport and metabolism | 196 | 22 | 10.09% |
| J | Translation, ribosomal structure and biogenesis | 208 | 26 | 11.11% |
| K | Transcription | 469 | 79 | 14.42% |
| L | Replication, recombination and repair | 144 | 26 | 15.29% |
| O | Post-translational modification, protein turnover, chaperones | 304 | 29 | 8.71% |
| P | Inorganic ion transport and metabolism | 243 | 23 | 8.64% |
| S | Function unknown | 1080 | 158 | 12.76% |
| T | Signal transduction mechanisms | 525 | 31 | 5.58% |

The data in Table 15 indicates areas for model enhancement. Misclassification rates in the ’K’ (Transcription) and ’T’ (Signal transduction mechanisms) categories suggest a need for improved feature engineering to better distinguish these proteins. Additionally, high misclassification in the ’S’ (Function unknown) category indicates a requirement for expanded training data and better annotations for proteins with unknown functions. Addressing these issues will help develop models that manage the complexities and overlaps in protein functions more effectively.

#### 4.5.3. Motif Analysis with MEME Suite

The motif analysis using MEME Suite identified patterns in motif occurrences between correctly classified and misclassified sequences across various tasks. The data presented in Table 16 provides insights into the occurrences of specific motifs in correctly classified and misclassified sequences across various protein-related tasks, including solubility, localization, ion channels, transporters, and membrane proteins (mp).

**Table 16.**
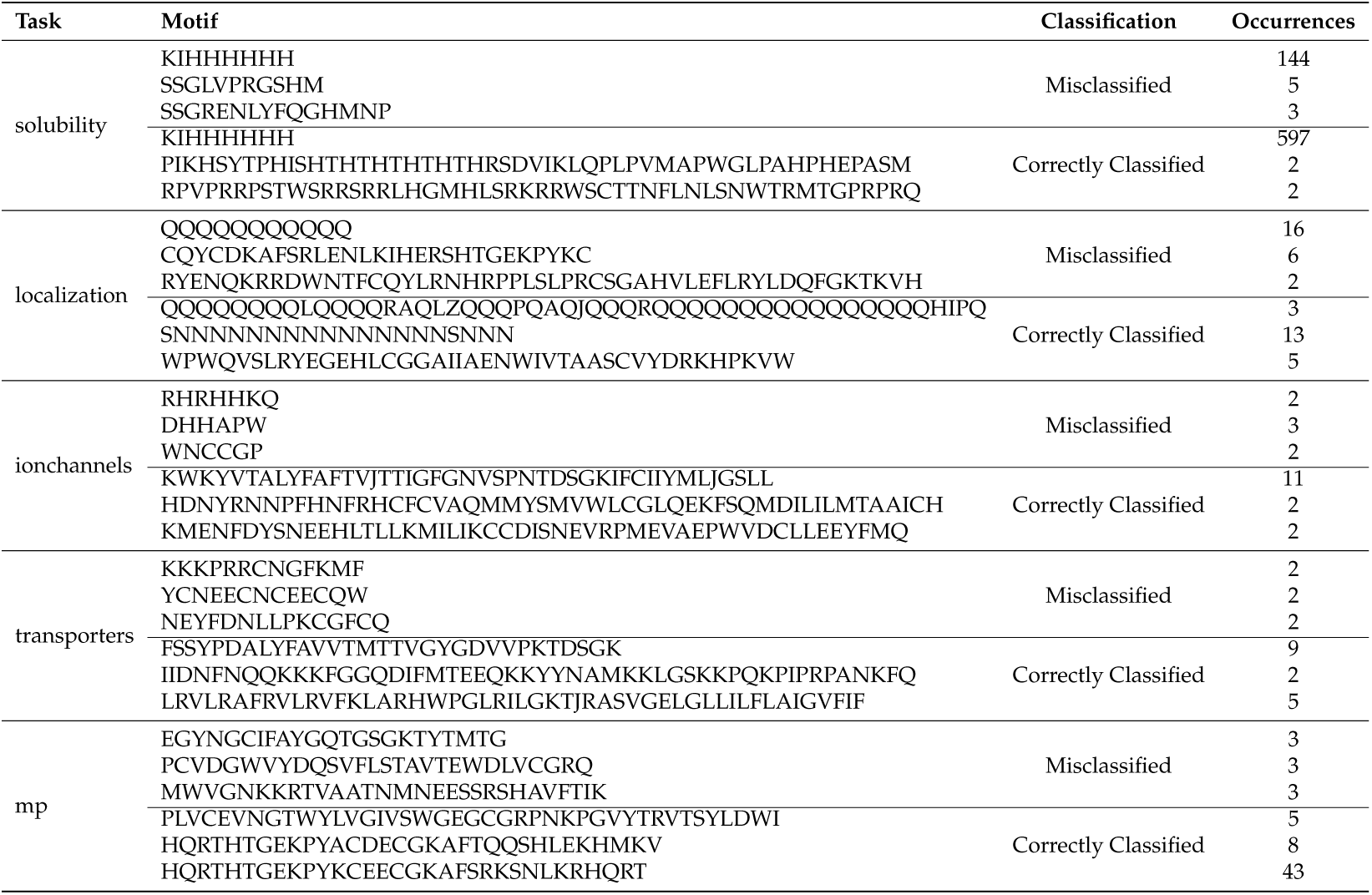
Motif occurrences in correctly and misclassified sequences. This table shows the complete list occurrences of specific motifs in sequences that were correctly classified and misclassified across various tasks. Each row details the task, the motif sequence, the classification status (whether the motif was found in correctly classified or misclassified sequences), and the number of occurrences of the motif. Note that “mp” stands for membrane proteins.

For the solubility task, the motif ’KIHHHHHH’ appeared 144 times in misclassified sequences compared to 597 times in correctly classified sequences, suggesting a potential correlation between this motif and correct solubility classification. The data indicates that this motif occurs significantly more often in sequences that were correctly classified as soluble, compared to those that were misclassified. This implies that the motif ’KIHHHHHH’ could be an important feature that the model uses to correctly identify solubility, and it might be contributing to the model’s ability to make accurate predictions regarding the solubility of proteins. However, this is an observation that would require further analysis to confirm its significance and to understand the underlying biological reasons for this correlation.

Other motifs such as ’SSGLVPRGSHM’ and ’SSGRENLYFQGHMNP’ were found less frequently in misclassified sequences, while motifs like ’PIKHSYTPHISHTHTHTHTHTHRS-DVIKLQPLPVMAPWGLPAHPHEPASM’ and ’RPVPRRPSTWSRRSRRLHGMHLSRKRRWSCTTNFLNLSNWTRMTGPRPRQ’ were rare in correctly classified sequences, indicating specific motifs’ varying impact on classification accuracy.

In the localization task, motifs such as ’QQQQQQQQQQQ’ and ’CQYCDKAFSRLEN-LKIHERSHTGEKPYKC’ were more frequent in misclassified sequences, with 16 and 6 occurrences respectively, while the motif ’SNNNNNNNNNNNNNNNSNNN’ appeared 13 times in correctly classified sequences. This pattern is consistent across other tasks, such as ion channels and transporters, where correctly classified sequences generally exhibited higher occurrences of certain motifs, like ’KWKYVTALYFAFTVJTTIGFGNVSP-NTDSGKIFCIIYMLJGSLL’ in ion channels (11 occurrences) and ’FSSYPDALYFAVVTMTTV-GYGDVVPKTDSGK’ in transporters (9 occurrences). The motif occurrences for membrane proteins also highlighted distinct patterns, with motifs such as ’HQRTHTGEKPYKCEECGKAF SRKSNLKRHQRT’ appearing 43 times in correctly classified sequences, indicating a significant motif presence correlating with accurate membrane protein classification.

## 5. Conclusion

This study aimed to enhance Protein Language Models (PLMs) by integrating secondary structure information, specifically alpha-helices and beta-sheets, into the TooT-PLM-P2S model. We evaluated TooT-PLM-P2S across several protein-related tasks, including fluorescence prediction, solubility prediction, sub-cellular localization, ion channel classification, transporter classification, membrane protein classification, and secondary structure prediction.

Our experiments demonstrated that incorporating secondary structure information into PLMs did not universally improve predictive accuracy. The Ankh model significantly outperformed TooT-PLM-P2S in three out of eight tasks. For the remaining five tasks, there were no statistically significant differences between the models, suggesting that any observed improvements in TooT-PLM-P2S could be due to random variations rather than true model enhancements. Notably, the only statistically significant improvement for TooT-PLM-P2S was in the precision metric for membrane protein classification, aligning with recent findings from other studies [52] when integrating 3D structural data.

These findings indicate that achieving statistical significance in future studies may require larger test sets or improved methods for integrating secondary structure information. Additionally, understanding how Ankh captures secondary structure information [53–55], particularly the roles of sequence identity and evolutionary conservation, is essential for refining these models. Future research should focus on expanding datasets and refining integration methods to enhance the robustness and accuracy of protein classification models.

## Author Contributions

Conceptualization, HG and GB; methodology, HG and GB; software, HG; validation, HG; formal analysis, HG and GB; investigation, HG; resources, HG and GB; data curation, HG; writing—original draft preparation, HG and GB; writing—review and editing, HG and GB; visualization, HG; supervision, GB; project administration, GB; All authors have read and agreed to the published version of the manuscript.

## Institutional Review Board Statement

Not applicable

## Informed Consent Statement

Not applicable

## Data Availability Statement

Not applicable

---

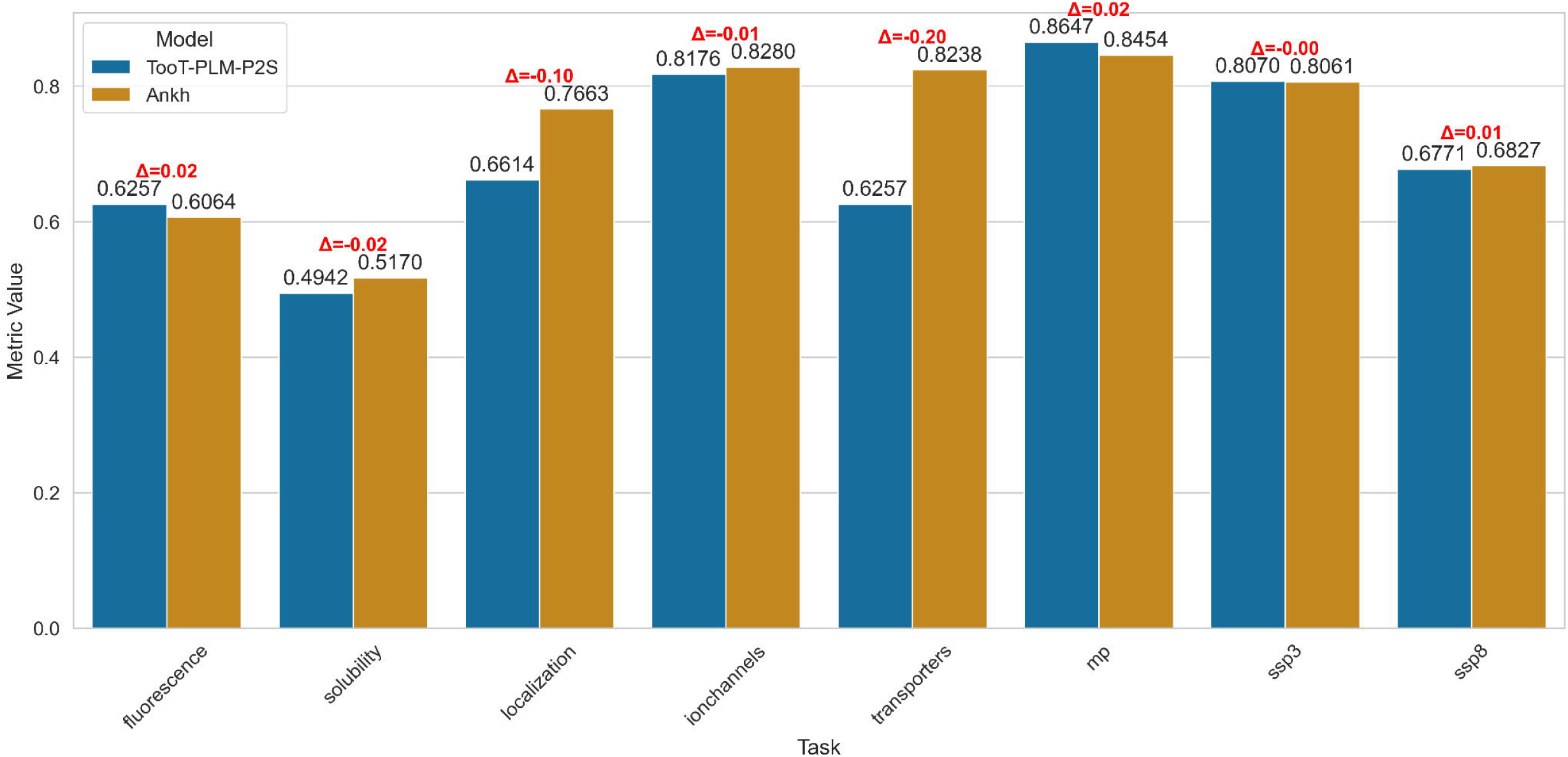

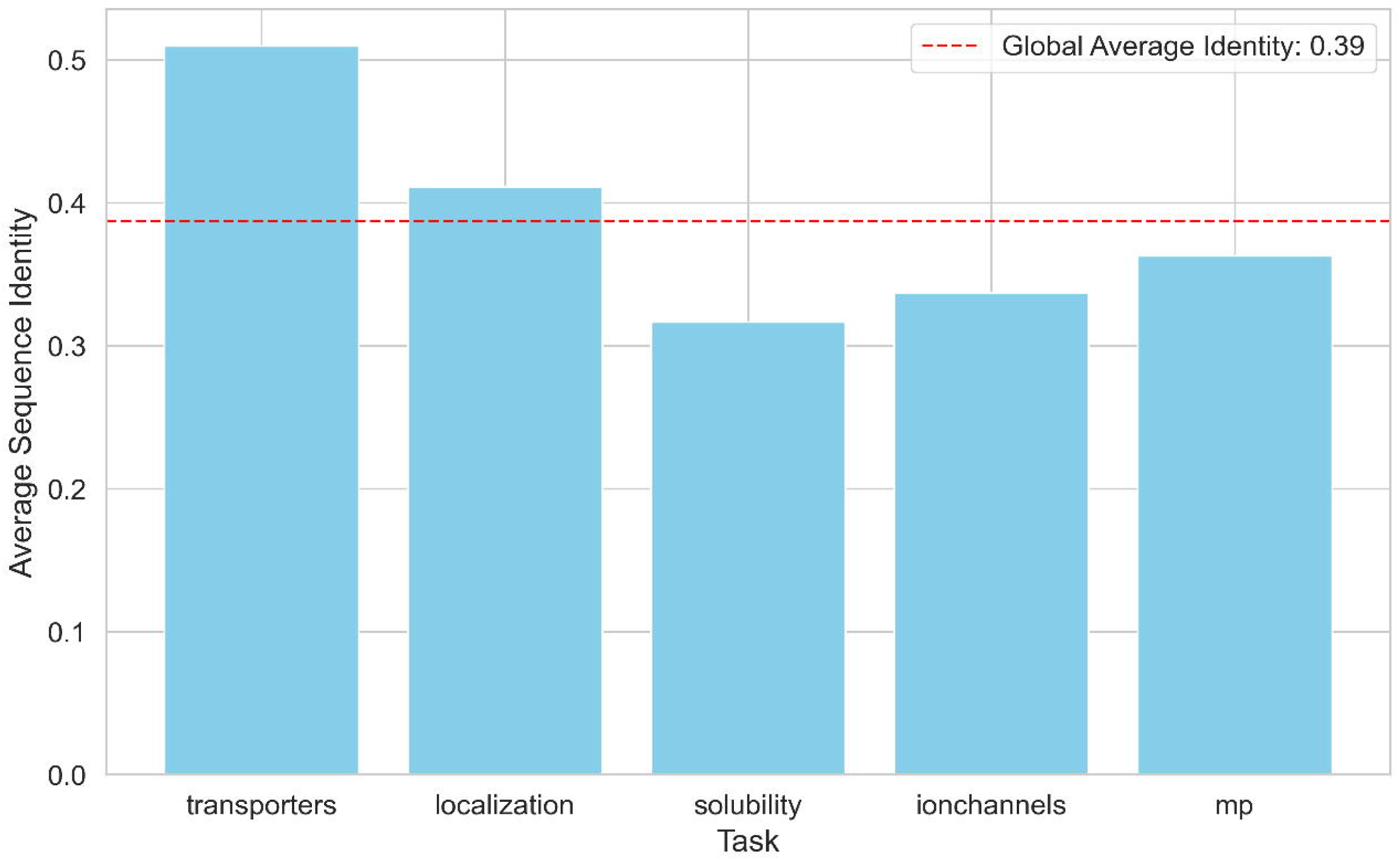

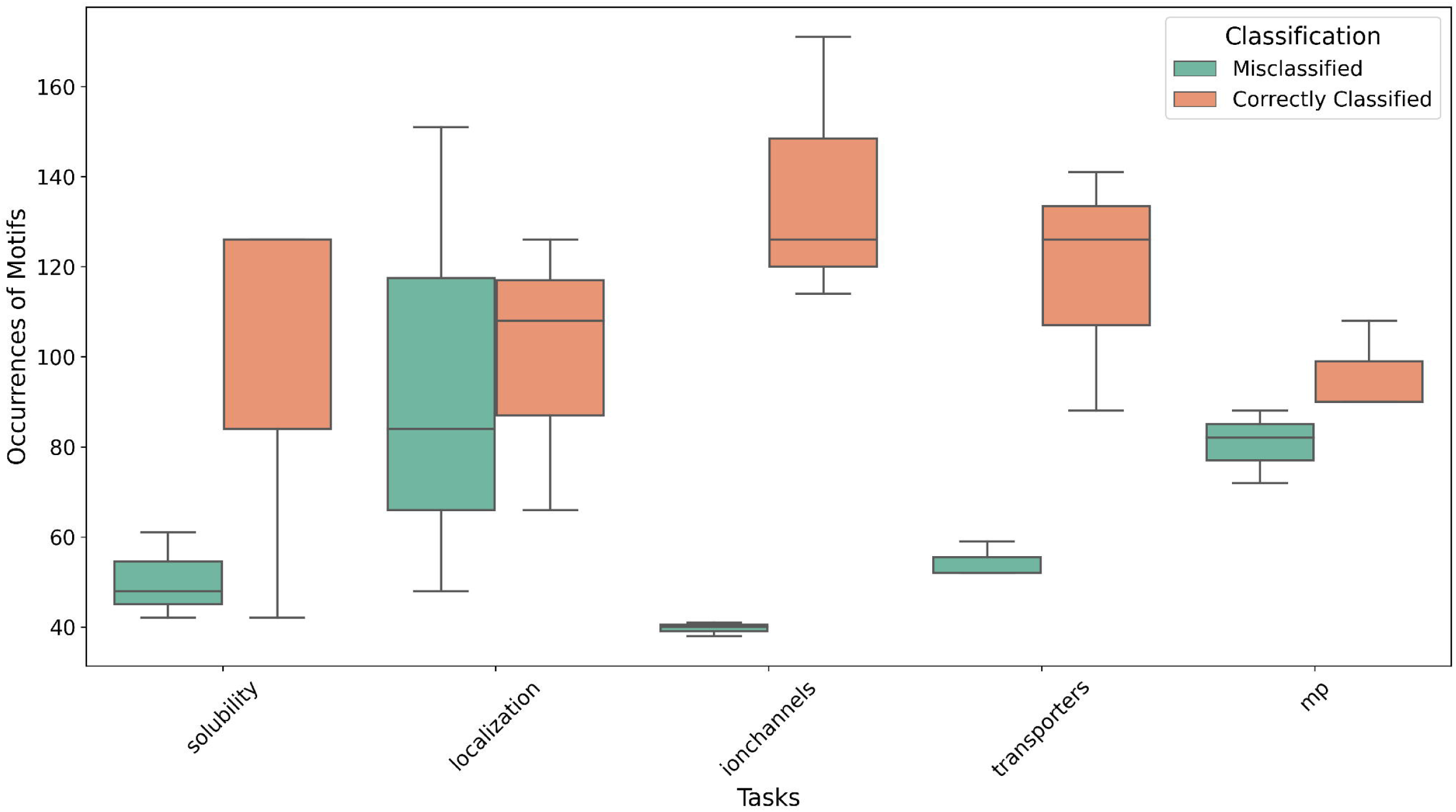

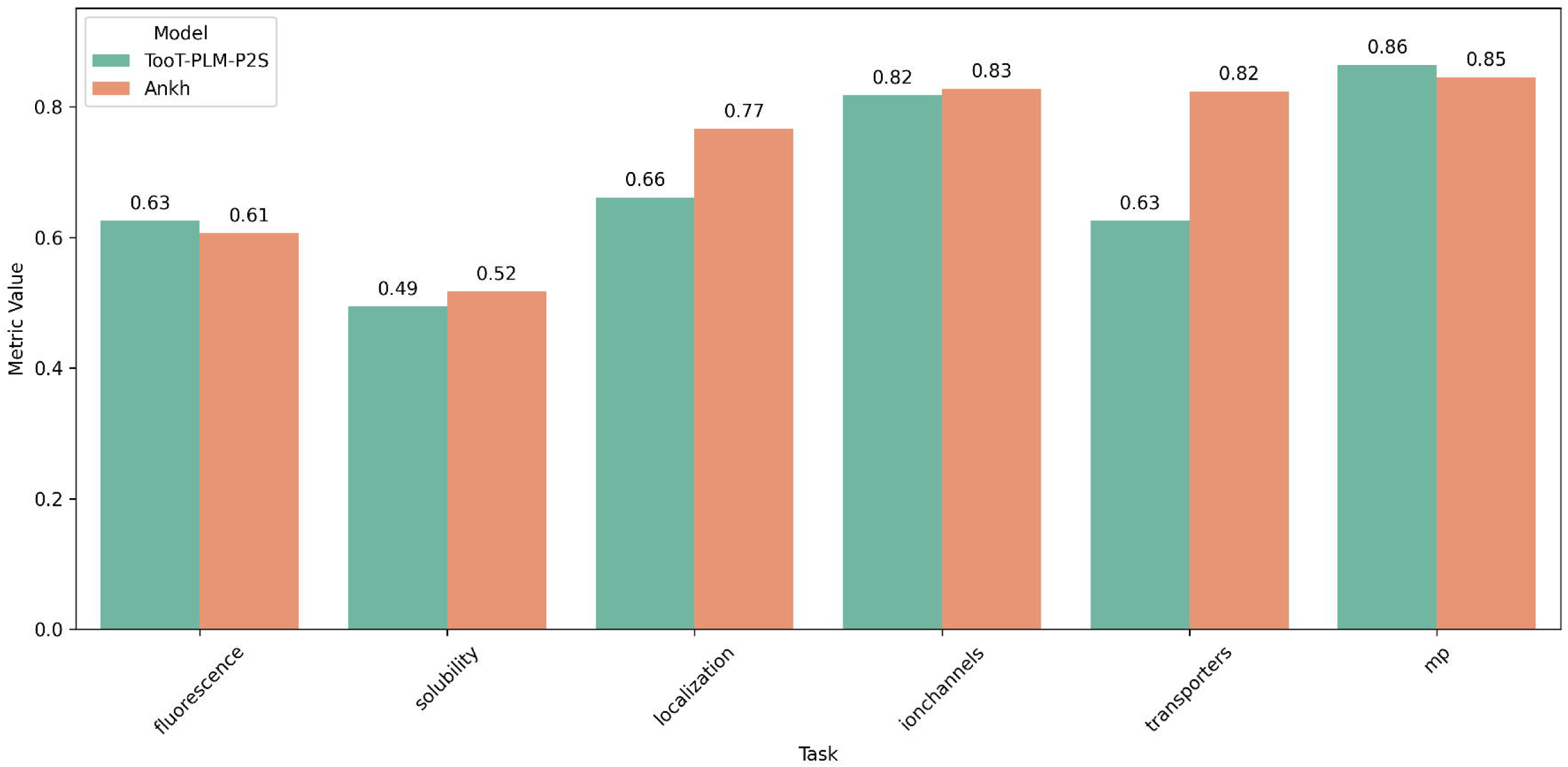

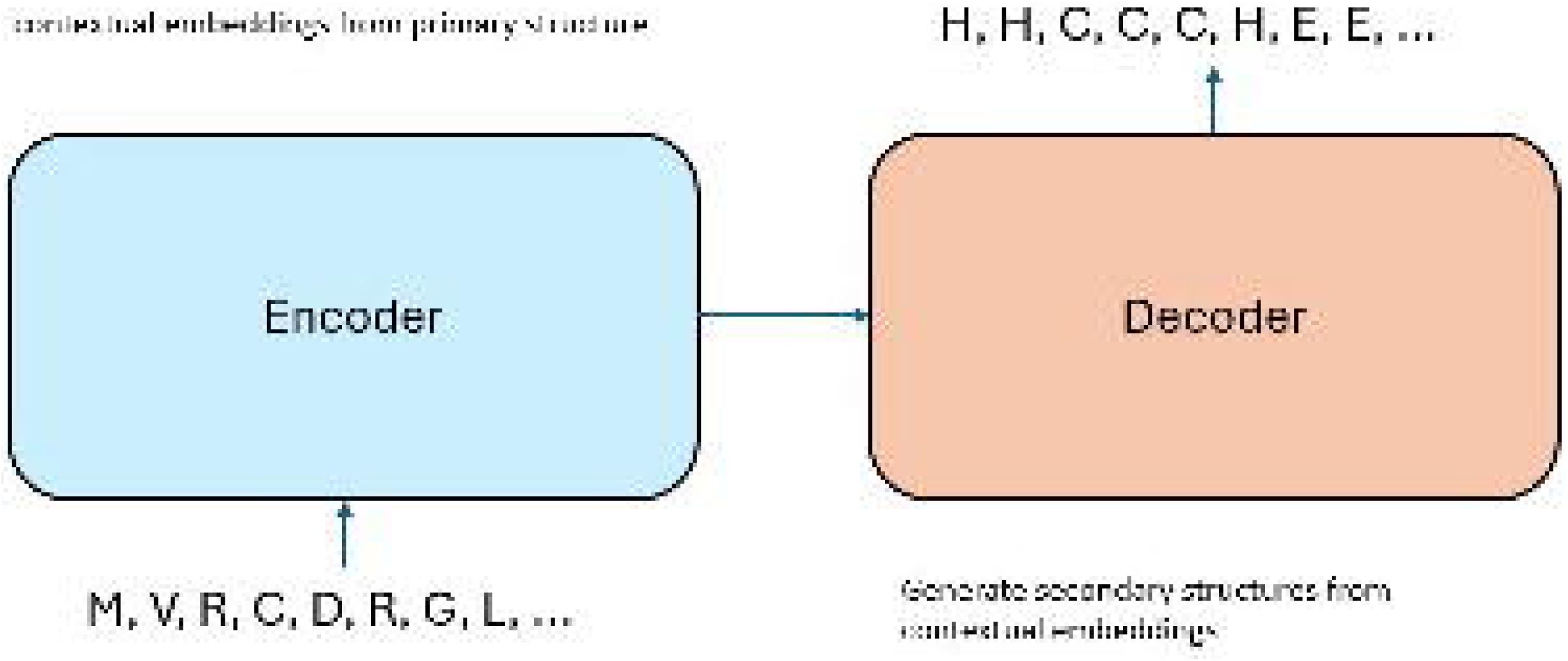

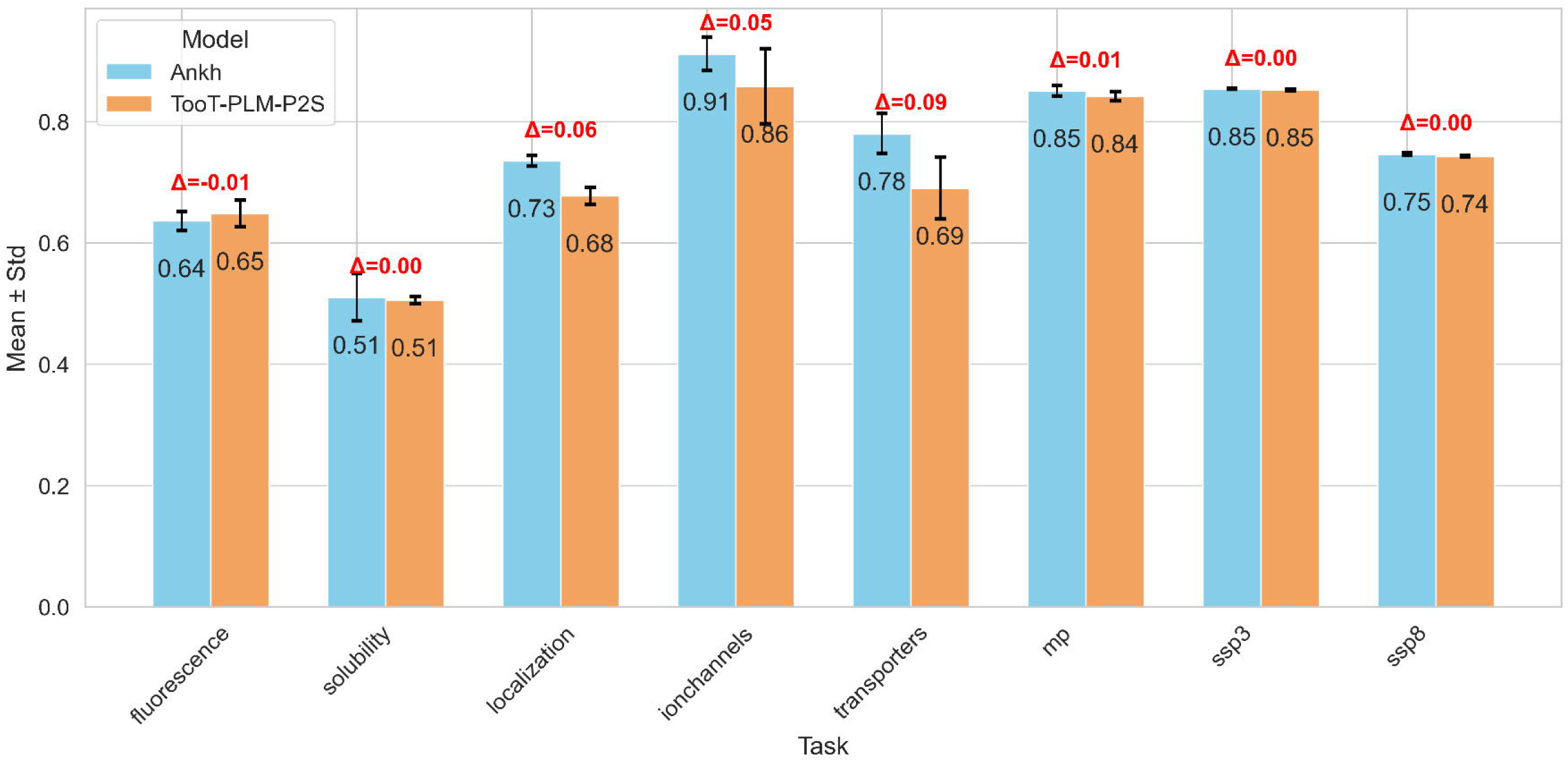

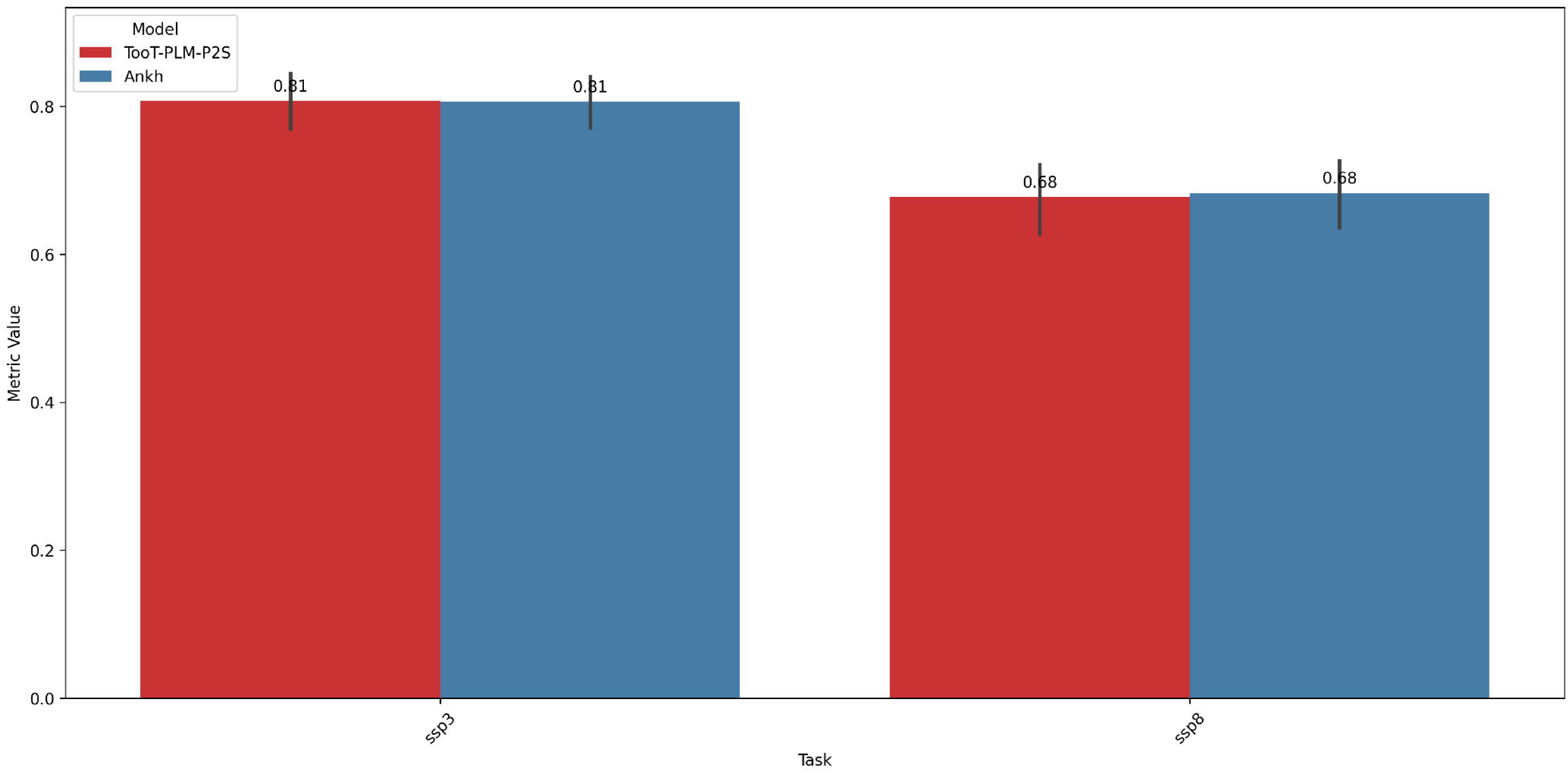

## References

1. Heinzinger, M.; Elnaggar, A.; Wang, Y.; Dallago, C.; Nechaev, D.; Matthes, F.; Rost, B. Modeling aspects of the language of life through transfer-learning protein sequences. BMC Bioinformatics 2019, 20, 723.

2. Rives, A.; Meier, J.; Sercu, T.; Goyal, S.; Lin, Z.; Liu, J.; Guo, D.; Ott, M.; Zitnick, C.L.; Ma, J.;, et al. Biological structure and function emerge from scaling unsupervised learning to 250 million protein sequences. Proceedings of the National Academy of Sciences 2021, 118, e2016239118. Publisher: Proceedings of the National Academy of Sciences.

3. Elnaggar, A.; Heinzinger, M.; Dallago, C.; Rehawi, G.; Wang, Y.; Jones, L.; Gibbs, T.; Feher, T.; Angerer, C.; Steinegger, M.;, et al. ProtTrans: Towards Cracking the Language of Lifes Code Through Self-Supervised Deep Learning and High Performance Computing. IEEE Transactions on Pattern Analysis and Machine Intelligence 2021, pp. 1–1.

4. Vaswani, A.; Shazeer, N.; Parmar, N.; Uszkoreit, J.; Jones, L.; Gomez, A.N.; Kaiser, L.; Polosukhin, I. Attention Is All You Need. *arXiv* 2017.

5. Liu, Y.; Ott, M.; Goyal, N.; Du, J.; Joshi, M.; Chen, D.; Levy, O.; Lewis, M.; Zettlemoyer, L.; Stoyanov, V. RoBERTa: A Robustly Optimized BERT Pretraining Approach, 2019. arXiv:1907.11692 [cs].

6. Raffel, C.; Shazeer, N.; Roberts, A.; Lee, K.; Narang, S.; Matena, M.; Zhou, Y.; Li, W.; Liu, P.J. Exploring the limits of transfer learning with a unified text-to-text Transformer, 2020. arXiv:1910.10683 [cs, stat].

7. Peters, M.E.; Neumann, M.; Iyyer, M.; Gardner, M.; Clark, C.; Lee, K.; Zettlemoyer, L. Deep contextualized word representations. In Proceedings of the Proceedings of the 2018 Conference of the North American Chapter of the Association for Computational Linguistics: Human Language Technologies, Volume 1 (Long Papers), New Orleans, Louisiana, 2018; pp. 2227–2237.

8. Asgari, E.; Mofrad, M.R.K. Continuous distributed representation of biological sequences for deep proteomics and genomics. PLOS ONE 2015, 10, e0141287.

9. Rao, R.M.; Liu, J.; Verkuil, R.; Meier, J.; Canny, J.; Abbeel, P.; Sercu, T.; Rives, A. MSA transformer. In Proceedings of the Proceedings of the 38th International Conference on Machine Learning. PMLR, 2021, pp. 8844–8856. ISSN: 2640-3498.

10. Rao, R.; Bhattacharya, N.; Thomas, N.; Duan, Y.; Chen, P.; Canny, J.; Abbeel, P.; Song, Y. Evaluating protein transfer learning with TAPE. In Proceedings of the Advances in Neural Information Processing Systems; Wallach, H.; Larochelle, H.; Beygelzimer, A.; Alché-Buc, F.d.; Fox, E.; Garnett, R., Eds. Curran Associates, Inc., 2019, Vol. 32.

11. Unsal, S.; Ataş, H.; Albayrak, M.; Turhan, K.; Acar, A.C.; Doğan, T. Evaluation of methods for protein representation learning: A quantitative analysis. Technical report, bioRxiv, 2020. Section: New Results Type: article.

12. Kotsiliti, E. De novo protein design with a language model. Nature Biotechnology 2022, 40, 1433–1433.

13. Ghazikhani, H.; Butler, G. A study on the application of protein language models in the analysis of membrane proteins. In Proceedings of the Distributed Computing and Artificial Intelligence, Special Sessions, 19th International Conference; Machado, J.M.; Chamoso, P.; Hernández, G.; Bocewicz, G.; Loukanova, R.; Jove, E.; del Rey, A.M.; Ricca, M., Eds., Cham, 2023; Lecture Notes in Networks and Systems, pp. 147–152.

14. Ghazikhani, H.; Butler, G. TooT-BERT-M: Discriminating membrane proteins from non-membrane proteins using a BERT representation of protein primary sequences. In Proceedings of the 2022 IEEE Conference on Computational Intelligence in Bioinformatics and Computational Biology (CIBCB), 2022, pp. 1–8.

15. Heinzinger, M.; Weissenow, K.; Sanchez, J.G.; Henkel, A.; Mirdita, M.; Steinegger, M.; Rost, B. Bilingual language model for protein sequence and structure, 2024. bioRxiv; Pages: 2023.07.23.550085 Section: New Results.

16. Wang, D.; Pourmirzaei, M.; Abbas, U.L.; Zeng, S.; Manshour, N.; Esmaili, F.; Poudel, B.; Jiang, Y.; Shao, Q.; Chen, J.; et al. S-PLM: Structure-aware protein language model via contrastive learning between sequence and structure, 2024. bioRxiv; Pages: 2023.08.06.552203 Section: New Results.

17. Varadi, M.; Bertoni, D.; Magana, P.; Paramval, U.; Pidruchna, I.; Radhakrishnan, M.; Tsenkov, M.; Nair, S.; Mirdita, M.; Yeo, J.;, et al. AlphaFold protein structure database in 2024: Providing structure coverage for over 214 million protein sequences. Nucleic Acids Research 2024, 52, D368–D375.

18. Ghazikhani, H.; Butler, G. Ion channel classification through machine learning and protein language model embeddings. Journal of Integrative Bioinformatics 2024. Under Review.

19. Elnaggar, A.; Essam, H.; Salah-Eldin, W.; Moustafa, W.; Elkerdawy, M.; Rochereau, C.; Rost, B. Ankh: Optimized protein language model unlocks general-purpose modelling, 2023. arXiv:2301.06568 [cs, q-bio].

20. Rehman, I.; Farooq, M.; Botelho, S. Biochemistry, Secondary Protein Structure. In StatPearls; StatPearls Publishing: Treasure Island (FL), 2022.

21. Buehler, L. The Structure of Membrane Proteins. In Cell Membranes; Garland Science, 2015. Section: 3.

22. Gromiha, M.M. Chapter 2 - Protein Sequence Analysis. In Protein Bioinformatics; Gromiha, M.M., Ed.; Academic Press: Singapore, 2010; pp. 29–62.

23. Quick, M.W., Ed. Transmembrane Transporters; Receptor biochemistry and methodology, Wiley-Liss: New York, 2002.

24. Tan, Y.; Li, M.; Zhou, B.; Zhong, B.; Zheng, L.; Tan, P.; Zhou, Z.; Yu, H.; Fan, G.; Hong, L. Simple, efficient and scalable structure-aware adapter boosts protein language models, 2024. arXiv:2404.14850 [cs, q-bio].

25. Su, J.; Han, C.; Zhou, Y.; Shan, J.; Zhou, X.; Yuan, F. SaProt: Protein language modeling with structure-aware vocabulary, 2024. bioRxiv; Pages: 2023.10.01.560349 Section: New Results.

26. Detlefsen, N.S.; Hauberg, S.; Boomsma, W. Learning meaningful representations of protein sequences. Nature Communications 2022, 13, 1914.

27. Stärk, H.; Dallago, C.; Heinzinger, M.; Rost, B. Light attention predicts protein location from the language of life. Bioinformatics Advances 2021, 1, vbab035.

28. Klausen, M.S.; Jespersen, M.C.; Nielsen, H.; Jensen, K.K.; Jurtz, V.I.; Sønderby, C.K.; Sommer, M.O.A.; Winther, O.; Nielsen, M.; Petersen, B.;, et al. NetSurfP-2.0: Improved prediction of protein structural features by integrated deep learning. Proteins: Structure, Function, and Bioinformatics 2019, 87, 520–527.

29. Xu, M.; Zhang, Z.; Lu, J.; Zhu, Z.; Zhang, Y.; Ma, C.; Liu, R.; Tang, J. PEER: A comprehensive and multi-task benchmark for protein sequence understanding, 2022. arXiv:2206.02096 [cs].

30. Chen, B.; Cheng, X.; Gengyang, L.a.; Li, S.; Zeng, X.; Wang, B.; Jing, G.; Liu, C.; Zeng, A.; Dong, Y.; et al. xTrimoPGLM: Unified 100B-scale pre-trained transformer for deciphering the language of protein, 2023. bioRxiv; Pages: 2023.07.05.547496 Section: New Results.

31. Sarkisyan, K.S.; Bolotin, D.A.; Meer, M.V.; Usmanova, D.R.; Mishin, A.S.; Sharonov, G.V.; Ivankov, D.N.; Bozhanova, N.G.; Baranov, M.S.; Soylemez, O.;, et al. Local fitness landscape of the green fluorescent protein. Nature 2016, 533, 397–401. Publisher: Nature Publishing Group.

32. Khurana, S.; Rawi, R.; Kunji, K.; Chuang, G.Y.; Bensmail, H.; Mall, R. DeepSol: a deep learning framework for sequence-based protein solubility prediction. Bioinformatics 2018, 34, 2605–2613.

33. Almagro Armenteros, J.J.; Sønderby, C.K.; Sønderby, S.K.; Nielsen, H.; Winther, O. DeepLoc: Prediction of protein subcellular localization using deep learning. Bioinformatics 2017, 33, 3387–3395.

34. Taju, S.W.; Ou, Y.Y. DeepIon: Deep learning approach for classifying ion transporters and ion channels from membrane proteins. Journal of Computational Chemistry 2019, 40, 1521–1529.

35. Mishra, N.K.; Chang, J.; Zhao, P.X. Prediction of Membrane Transport Proteins and Their Substrate Specificities Using Primary Sequence Information. PLOS ONE 2014, 9, e100278.

36. Alballa, M.; Butler, G. Integrative approach for detecting membrane proteins. BMC Bioinformatics 2020, 21, 575.

37. Yang, Y.; Gao, J.; Wang, J.; Heffernan, R.; Hanson, J.; Paliwal, K.; Zhou, Y. Sixty-five years of the long march in protein secondary structure prediction: the final stretch? Briefings in Bioinformatics 2018, 19, 482–494.

38. Cuff, J.A.; Barton, G.J. Evaluation and improvement of multiple sequence methods for protein secondary structure prediction. *Proteins: Structure*, Function, and Bioinformatics 1999, 34, 508–519.

39. Abriata, L.A.; Tamò, G.E.; Monastyrskyy, B.; Kryshtafovych, A.; Dal Peraro, M. Assessment of hard target modeling in CASP12 reveals an emerging role of alignment-based contact prediction methods. *Proteins: Structure*, Function, and Bioinformatics 2018, 86, 97–112.

40. Kryshtafovych, A.; Schwede, T.; Topf, M.; Fidelis, K.; Moult, J. Critical assessment of methods of protein structure prediction (CASP)—Round XIV. *Proteins: Structure*, Function, and Bioinformatics 2021, 89, 1607–1617.

41. Jiang, Z.H.; Yu, W.; Zhou, D.; Chen, Y.; Feng, J.; Yan, S. ConvBERT: Improving BERT with span-based dynamic convolution. In Proceedings of the Advances in Neural Information Processing Systems. Curran Associates, Inc., 2020, Vol. 33, pp. 12837–12848.

42. Hendrycks, D.; Gimpel, K. Gaussian error linear units (GELUs), 2023. arXiv:1606.08415 [cs].

43. Akiba, T.; Sano, S.; Yanase, T.; Ohta, T.; Koyama, M. Optuna: A next-generation hyperparameter optimization framework. In Proceedings of the Proceedings of the 25th ACM SIGKDD International Conference on Knowledge Discovery & Data Mining, New York, NY, USA, 2019; KDD ’19, pp. 2623–2631.

44. Chicco, D.; Jurman, G. The advantages of the Matthews Correlation Coefficient (MCC) over F1 score and accuracy in binary classification evaluation. BMC Genomics 2020, 21, 6.

45. Mowery, B.D. The paired t-test. Pediatric Nursing 2011, 37, 320–322. Publisher: Jannetti Publications, Inc.

46. Subramanian, A.; Tamayo, P.; Mootha, V.K.; Mukherjee, S.; Ebert, B.L.; Gillette, M.A.; Paulovich, A.; Pomeroy, S.L.; Golub, T.R.; Lander, E.S.;, et al. Gene set enrichment analysis: A knowledge-based approach for interpreting genome-wide expression profiles. Proceedings of the National Academy of Sciences 2005, 102, 15545–15550. Publisher: Proceedings of the National Academy of Sciences.

47. Reimand, J.; Isserlin, R.; Voisin, V.; Kucera, M.; Tannus-Lopes, C.; Rostamianfar, A.; Wadi, L.; Meyer, M.; Wong, J.; Xu, C.;, et al. Pathway enrichment analysis and visualization of omics data using g:Profiler, GSEA, Cytoscape and EnrichmentMap. Nature Protocols 2019, 14, 482–517. Publisher: Nature Publishing Group.

48. Notredame, C.; Higgins, D.G.; Heringa, J. T-coffee: A novel method for fast and accurate multiple sequence alignment. Journal of Molecular Biology 2000, 302, 205–217.

49. Cock, P.J.A.; Antao, T.; Chang, J.T.; Chapman, B.A.; Cox, C.J.; Dalke, A.; Friedberg, I.; Hamelryck, T.; Kauff, F.; Wilczynski, B.;, et al. Biopython: Freely available Python tools for computational molecular biology and bioinformatics. Bioinformatics 2009, 25, 1422–1423.

50. Huerta-Cepas, J.; Szklarczyk, D.; Heller, D.; Hernández-Plaza, A.; Forslund, S.K.; Cook, H.; Mende, D.R.; Letunic, I.; Rattei, T.; Jensen, L.;, et al. eggNOG 5.0: a hierarchical, functionally and phylogenetically annotated orthology resource based on 5090 organisms and 2502 viruses. Nucleic Acids Research 2019, 47, D309–D314.

51. Bailey, T.L.; Johnson, J.; Grant, C.E.; Noble, W.S. The MEME Suite. Nucleic Acids Research 2015, 43, W39–W49.

52. Heinzinger, M.; Weissenow, K.; Sanchez, J.G.; Henkel, A.; Steinegger, M.; Rost, B. ProstT5: Bilingual language model for protein sequence and structure. preprint, Bioinformatics, 2023.

53. Chowdhury, R.; Bouatta, N.; Biswas, S.; Floristean, C.; Kharkar, A.; Roy, K.; Rochereau, C.; Ahdritz, G.; Zhang, J.; Church, G.M.;, et al. Single-sequence protein structure prediction using a language model and deep learning. Nature Biotechnology 2022, 40, 1617–1623. Publisher: Nature Publishing Group.

54. Lin, Z.; Akin, H.; Rao, R.; Hie, B.; Zhu, Z.; Lu, W.; Costa, A.d.S.; Fazel-Zarandi, M.; Sercu, T.; Candido, S.; et al. Language models of protein sequences at the scale of evolution enable accurate structure prediction, 2022. bioRxiv; Pages: 2022.07.20.500902 Section: New Results.

55. Vig, J.; Madani, A.; Varshney, L.R.; Xiong, C.; Socher, R.; Rajani, N.F. BERTology Meets Biology: interpreting attention in protein language models, 2021. arXiv:2006.15222 [cs, q-bio].

